# Effective *in vivo* RNA base editing *via* engineered cytidine deaminase APOBECs fused with PUF proteins

**DOI:** 10.1101/2025.02.04.636546

**Authors:** Wenjian Han, Bo Yuan, Xiaojuan Fan, Weike Li, Yiting Yuan, Yuefang Zhang, Shu Wang, Shifang Shan, Markus Hafner, Zefeng Wang, Zilong Qiu

## Abstract

Base editing stands at the forefront of genetic engineering, heralding precise genetic modifications with broad implications. While CRISPR-based DNA and RNA base editing systems capitalize on sgRNA-guided specificity and diverse deaminase functionalities, the pursuit of efficient C-to-U RNA editing has been hampered by the inherent constraints of cytidine deaminases. Here, we report an RNA base editing platform by refining cytidine deaminases, termed professional APOBECs (ProAPOBECs), through systematic enhancements and AI-driven protein engineering. ProAPOBECs demonstrate unprecedented catalytic versatility, particularly fused with RNA-recognizing Pumilio and FBF (PUF) proteins. We present the first effective use of RNA base editing in the brain with ProAPOBECs in Mef2c mutant mice, a model for autism. The AAV-mediated RNA base editing via ProAPOBEC not only corrects genetic mutation in mRNAs but significantly alleviates autistic-like behaviors in the mice. This work introduces a pioneering collection of RNA base editing instruments, emphasizing their therapeutic potential in combatting genetic disorders.

## Introduction

CRISPR-based base editing offers a potent method for editing genetic material by directly modifying single base pair in DNA or RNA. This approach, involving C-to-U (cytidine-to-uridine) and A-to-I (adenosine-to-inosine) transitions mediated by deaminases, avoids the strand breaks typically induced by the CRISPR/Cas system ^1,2^. Initially developed for DNA modification, CRISPR/Cas-based targeting has become an effective tool in genetic research and holds potential for therapeutic applications, such as correcting disease-associated mutations in animal models ^3–9^.

Unlike the permanent modifications made by DNA base editing ^10–12^, RNA base editing offers a reversible method for *in vivo* base alternations, providing a relatively safer avenue for therapeutic strategies ^13–17^. The A-to-I transition in RNA, mediated by Adenosine Deaminases Acting on RNA (ADAR), has been effectively demonstrated *in vivo* ^18–20^. Two evolved ADAR2-based editing systems, RESCUE-S and xCBE, have been developed to achieve both A-to-I and C-to-U RNA editing ^21,22^. However, RESCUE-S is strongly biased against editing G<u>C</u> and C<u>C</u> motifs, limiting its versatility ^23^. To address this limitation, Stafforst’s group introduced the SNAP-CDAR-S system, a SNAP-tag-based tool designed to edit C<u>C</u> motifs in specific RNA targets ^24^. While promising, evolved ADAR2-mediated base editing systems, including SNAP-CDAR-S, exhibit simultaneous C-to-U and A-to-I off-target editing, imposing constraints on their broader application ^21–24^. Engineering Apolipoprotein B mRNA editing enzyme catalytic polypeptide (APOBEC) family proteins offers a potential solution. These proteins are specifically tailored for precise C-to-U base editing, effectively eliminating the A-to-I off-target effects and reducing undesired side effects. This approach represents a significant advancement in RNA base editing technology.

More recently, an RNA base editing system called CURE has been developed utilizing the dCasRx protein and a native APOBEC3A, to achieve limited yet specific U<u>C</u> transitions ^23^. However, the specific Cas13-based C-to-U transition, has not achieved for efficiently correcting disease-causing mutations in mouse models ^23,25^. One potential reason is the strong preference of APOBEC for single-stranded RNA, conflicting with the double-stranded structures formed by target mRNA and small guide RNA used in the CRISPR/Cas system ^26^.

To address existing limitations, the gRNA-free system, REWIRE (RNA editing with individual RNA-binding enzyme), was developed by exploiting the programmable RNA targeting capabilities of PUF proteins ^27^. PUF proteins feature 8- or 10- repeat motifs, each of which can be programmed to specifically bind any RNA base *via* interaction with the Watson–Crick edge ^28–32^. Various groups have developed PUF-based systems incorporating different functional domains to manipulate RNA splicing ^33^, translation ^34^, degradation ^35^ and methylation ^36^. Given the versatility of PUF domain in recognizing nearly any short 8- or 10- nucleotide RNA sequences, its combination with ADAR or APOBEC enzymes allows REWIRE to perform precise and efficient A- to-I or C-to-U editing on specific nucleosides of RNA targets in cultured cells ^27^. Notably, the use of mammalian-derived proteins in the REWIRE minimizes the potential immune responses often triggered by bacterial Cas enzymes in therapeutic base editing scenarios ^37^.

Current C-to-U RNA base editing systems rely on the enzymatic activity of APOBEC to hydrolytically deaminate cytidine ^13,27^. Natural APOBECs, however, often exhibit deamination activity in a sequence context-dependent manner, primarily with single-strand DNA or RNA ^38^. For instance, Human APOBEC3A preferentially edits cytidine within UC motifs in mRNA ^23,27^. These context preferences limited further application of RNA base editors.

APOBEC enzymes, recognized for their critical role in inducing mutations within the genomes of retroviruses during infection, constitute a significant group of antiviral genes. Current NCBI database has documented over 8,231 eukaryotic genes homologous to the AID/APOBEC family, highlighting their extensive presence and diversity in eukaryotic organisms ^39^. These AID/APOBEC proteins share a common domain configuration, with a highly conserved catalytic domain centered between the variable N-terminal (NTD) and C-terminal domain (CTD) ^39–41^, which may enable a rational design of novel cytidine deaminases using artificial intelligence-assisted protein engineering.

In this study, we use AlphaFold2-mediated structural engineering to develop Professional APOBECs (ProAPOBECs) with greatly expanded C-to-U editing capability. When integrated within the REWIRE system, ProAPOBECs demonstrate improved specificity and efficiency in multiple sequence contexts, including G<u>C</u>, C<u>C</u>, A<u>C</u>, and U<u>C</u>.

Importantly, we achieved effective *in vivo* C-to-U RNA editing in the liver and brain of mice using the CU-REWIRE5s with ProAPOBECs delivered *via* adeno-associated virus (AAV). This *in vivo* RNA editing mediated by CU-REWIRE successfully alleviated autistic-like behaviors by correcting a point mutation in Mef2c mRNA in an autism spectrum disorder (ASD) mouse model ^8^. In conclusion, this study demonstrated that the AI-assisted protein engineering can be applied to refine the activity of RNA base editors, enabling the correction of autistic-like phenotype in mouse model and paving the way for the practical applications of RNA base editing in the realm of gene therapy.

## Results

### Enhanced Stability and Efficacy of CU-REWIRE *via* Structural Optimization of PUF Domain

Our initial efforts focused on improving CU-REWIRE3.0, which combined a 10-repeat PUF domain (PUF10) with the cytidine deaminase enzyme APOBEC3A. Structural analysis revealed high similarity between the fourth repeat (R4) of PUF10 in CU-REWIRE3.0 and the fifth repeat (R5) of the native Pumilio 2 protein. Importantly, R4 lacked the Leucine-Proline (LP) peptide present in R5, a potentially crucial element for structural flexibility as suggested by the previous research ^42^. This insight led us to engineer an enhanced PUF10 (ePUF10) with the LP peptide integrated into R4, resulting in the development of CU-REWIRE4.0 (Figures 1A and S1A).

**Figure 1.**
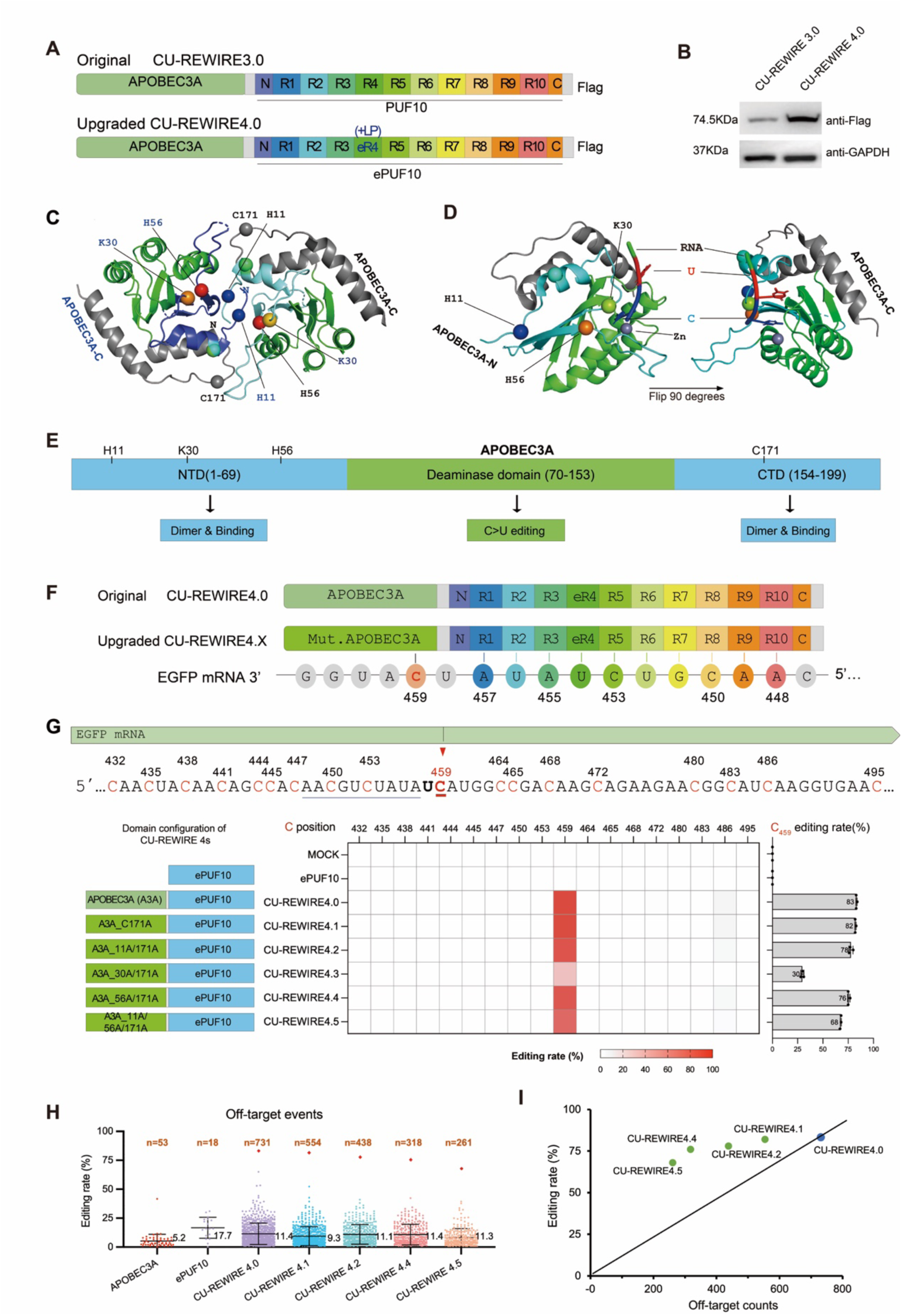
Structure-based optimization of APOBEC3A enhances editing efficacy and reduces off-target effects of REWIRE-mediated RNA base editing. (A) Schematic design of REWIREs with PUF variants. Top: Original CU-REWIRE 3.0 with PUF10, targeting 10-nt sequences. Bottom: Upgraded CU-REWIRE 4.0 with ePUF10. (B) Expression levels of CU-REWIREs in cultured cells. Representative Western blot images using anti-Flag antibody, related to (Figure S1B). (C) Crystal structure of Human APOBEC3A (PDB: 4XXO) highlighting putative dimerization interactions. Amino acids involved in dimer formation are shown in spherical forms. (D) Crystal structure of Human APOBEC3A (PDB: 5KEG) in complex with single-stranded DNA. Amino acids responsible for RNA recognition are labeled in spherical forms. Zn symbolizes the catalytic core. (E) Schematic illustration of APOBEC3A with its functional domains. (F) Schematic showing interaction between target EGFP mRNA and CU-REWIRE4.0 and 4.X with modified APOBEC3A variants. (G) Base editing efficiency of EGFP mRNA by CU-REWIRE4.X. The PUF binding site in EGFP is underlined in blue, with adjacent cytosines marked in red (top). Heatmap displays editing rates of cytosines near the on-target site C_459_ for each CU-REWIRE and control (ePUF10 alone), with C_459_ editing rate detailed on the right. Editing rates were measured by RNA-seq with triplicates (refer to methods). (H) Global editing rates of transcriptome-wide RNA base editing in samples treated with different CU-REWIRE4.X (related to data from panel g, labels and cutoff as in Supplementary Fig. 1e). Orange rhombuses indicate the on-target site EGFP-C_459_, and the mean number of off-target editing events is noted beside the dot plots. n represents the total number of edited cytosines detected. (I) Correlation between on-target editing rates (Y-axis) and transcriptome-wide C-to-U off-target editing events (X-axis) for various CU-REWIRE4s.

Comparative analysis showed that CU-REWIRE4.0 exhibited higher expression levels than its predecessor, suggesting improved stability due to the LP insertion (Figure 1B). We then assessed the editing efficacy of CU-REWIRE4.0 by targeting the C-to-U transition on the C_459_ site in the Enhanced Green Fluorescent Protein (EGFP) mRNA. As judged by mRNA sequencing (mRNA-seq), the CU-REWIRE4.0 achieved a substantial increase in editing efficiency, with an 82.3% success rate compared to 69.7% for CU-REWIRE3.0 (Figure S1B).

To evaluate off-target effects, we conducted RNA-seq analysis with 50X transcriptome coverage (See Methods for details). Four independent sample sets (APOBEC3A, ePUF10, CU-REWIRE3.0, and CU-REWIRE4.0) were analyzed with mRNA-seq, revealing 224 potential off-target events for CU-REWIRE3.0 and 731 for CU-REWIRE4.0 (Figures S1C-S1E). Notably, none of the off-target sites were within 20-nt downstream of ePUF10-binding sequences (Figure S1E), suggesting a high precision of ePUF10 in recognizing target and a significantly lower off-target rate compared to the earlier CU-REWIREs ^27^. However, off-target events were largely attributed to the basal activity of APOBECs, highlighting the need for further optimization.

While showing promise in editing efficiency, the original CU-REWIRE3.0 was restricted to C-to-U editing activity within the UC context ^27^. Similar to its predecessors, we found that the CU-REWIRE4.0 maintained the preference for editing cytidines within the UC consensus motif (Figure S1F). Its precision was further improved in that the C-to-U conversions predominantly occurred at the second position downstream of the ePUF10 binding site (Figure S1G). This underscores the enhanced specificity and efficiency of CU-REWIRE4.0.

### Structure-Based Optimization of APOBEC3A Minimizes Off-Target Effects

APOBEC3A proteins tend to form dimers that stabilize their interaction with single-stranded DNA or RNA ^38^. While this interaction is key in inducing mutations in viral genomes, it also contributes to bystander editing and off-target effects in non-target genomic regions during base editing. Therefore, our next objective was to reduce APOBEC3A dimerization.

By investigating the crystalline structure of APOBEC3A and building upon previous research, we identified two amino acid sites (H11, C171) critical for dimerization ^38^. The mutation of these sites could reduce the dimer formation and the association with nucleic acids of APOBEC3A (Figure 1C). Additionally, two amino acids (K30, H56) were found to be involved in substrate RNA recognition by APOBEC3A ^38^. Thus we speculate that mutations at these sites may diminish RNA recognition and, consequently, reduce off-target effects (Figure 1D).

To test this possibility, we introduced various point mutations into human APOBEC3A (Figure 1E). We then fused these mutated APOBEC3A versions with the ePUF10 domain, creating multiple CU-REWIRE4 variants that target C_459_ of EGFP mRNA expressed in HEK293T cells (Figure 1F). After co-transfecting these constructs with the EGFP expressing plasmid, we evaluated the C-to-U editing efficiency at the C_459_ site of EGFP mRNA by these CU-REWIRE4 variants (Figure 1G).

One variant, CU-REWIRE 4.1, with a C171A substitution in the C-terminal domain (CTD) of APOBEC3A, showed high editing efficiency while significantly reducing off-target effects (Figures 1G and 1H). Interestingly, the CU-REWIRE 4.2, 4.4, and 4.5, each containing point mutations in the NTD, showed moderate editing efficiency but substantially minimized off-target events. These findings indicate that strategically designed point mutations in the NTD and CTD of APOBEC3A can significantly decrease off-target effects while maintaining sufficient editing efficiency (Figure 1H). We further examined the on-target editing rates and off-target editing events of each CU-REWIRE variant, and found that these newly engineered variants provide optimal ratios of on- *vs.* off-target editing rates, which may offer versatile options for base editing applications (Figure1I).

Like the original CU-REWIREs, the modified CU-REWIRE4 variants predominantly edited cytidines within the UC motif (Figure S2A), reflecting the intrinsic target preference of APOBEC3A ^13,27^. To broaden the application scope of RNA base editing, we explored whether the APOBEC-derived base editors can achieve C-to-U editing in other context (i.e., C<u>C</u>, G<u>C</u>, and A<u>C</u> motifs) by systematically screening different AID/APOBEC enzymes that are integrated in the CU-REWIRE platform (Figure S2B). We found that only the APOBEC3A-ePUF10 configuration showed detectable base editing near the ePUF10 binding site (Figure S2C), and therefore select APOBEC3A as the chassis to introduce additional modifications that may expand the editing capabilities of new generation CU-REWIREs.

### AI-Assisted Engineering of Professional APOBECs (ProAPOBECs)

Our analysis of AID/APOBEC proteins revealed that while the deaminase domain is relatively conserved, the NTD and CTD are prone to evolutionary changes ^39–41^. This implies that the deaminase domain is crucial for catalyzing the C-to-U transition, while the NTD and CTD may specialize in recognizing different DNA/RNA targets ^39^. Consistent with this notion, previous studies suggested that various APOBEC proteins might exhibit C-to-U activity outside of the U<u>C</u> context ^13^, leading us to hypothesize that the core deaminase domain of APOBEC proteins may potentially edit cytosine in diverse contexts when combined with distinct NTD or CTD domains.

To test this hypothesis, we utilized AlphaFold2 to analyze the structures of APOBEC3 and APOBEC1 from both human and rodent origins (Figure 2A), similar to a previously used approach that analyzes deaminases across different species ^43^. These two enzymes are the only cytosine deaminases that are currently effective as active domains in DNA and RNA base editors ^13^. We analyzed the phylogenetic tree of AID/APOBEC family using 1160 full length eukaryotic AID/APOBEC-related genes documented in the NCBI database, and found that the APOBEC3 and APOBEC1 belong to the closely related clade of this tree (Figure 2B). There is striking similarity in the core deaminase domains of APOBEC3s and APOBEC1s, suggesting that functional modules may be interchangeable among these proteins. Therefore we combined the core deaminase domain of different APOBECs with the NTD and CTD of human APOBEC3A. These hybrid deaminases were termed ProAPOBECs, potentially applicable for both DNA and RNA base editing (Figure 2C). Together, over thousands of functional ProAPOBEC proteins could be predicted (Figure 2D).

**Figure 2.**
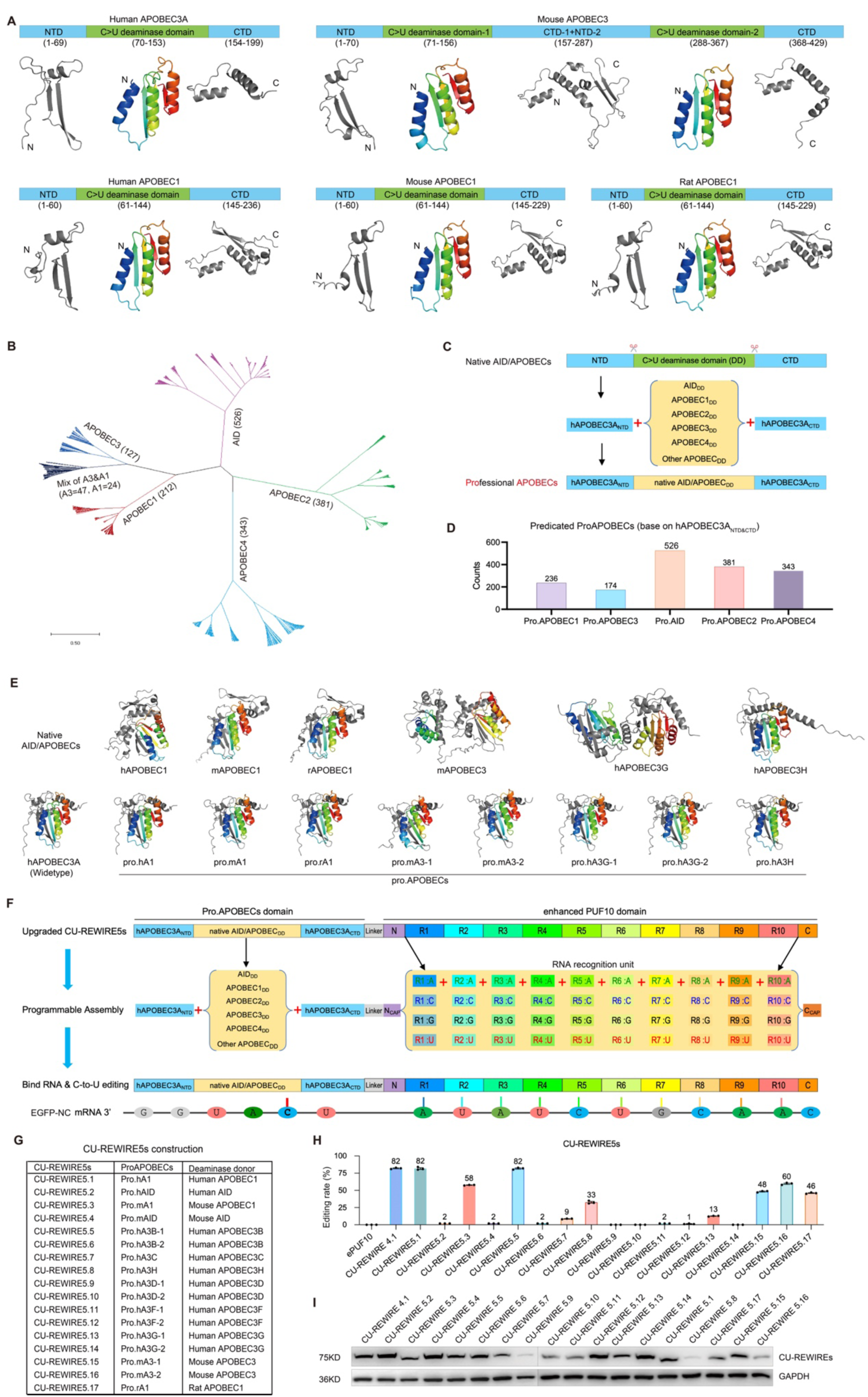
AI-Assisted design of programmable APOBEC platform for engineering ProAPOBEC enzymes. (A) Crystal structure of Human APOBEC3A (PDB: 4XXO), AlphaFold2-predicted structures of native APOBEC proteins: Mouse APOBEC3, Human APOBEC1, Mouse APOBEC1, and Rat APOBEC1, showcasing the NTD, C-to-U deaminase domain, and CTD. (B) Phylogenetic tree of the AID/APOBEC family, with major clades highlighted in different colors. (C) Schematic representation of the assembly of ProAPOBECs. (D) Estimated number of potential ProAPOBEC candidates based on AI-predicted AID/APOBEC deaminase domains. (E) AlphaFold2-predicted structures of AID/APOBEC proteins. Top: AI-predicted structures of native AID/APOBECs. Bottom: AI-predicted structures of ProAPOBECs. (F) The programmable CU-REWIRE5 system with ProAPOBEC variants. Top: Upgraded CU-REWIRE5, featuring integration of ProAPOBEC variants for targeted C-to-U editing. Middle: Schematic depiction of the modular assembly process of CU-REWIRE5. The ProAPOBECs domain, constructed from a selection of natural APOBEC deaminase domains (APOBEC_DD_) derived from various sources, is engineered for specific C-to-U base editing. Accompanying this, the ePUF10 domain includes 10 RNA recognition units, each designed for the specific detection of a unique RNA base type. Bottom: Illustration of the precise engagement of CU-REWIRE5 with the target RNA, showcasing the accurately binding and editing C_459_ of EGFP mRNA. (G) Illustrations depicting the components of CU-REWIRE5, ProAPOBECs, and their deaminase donors. (H) Various editing rates on C_459_ of EGFP mRNA of CU-REWIRE5s, assembled with ProAPOBECs in panel (i) and Supplementary Fig. 2d.(I) Protein expression levels of CU-REWIRE5s in HEK293T cells. The CU-REWIREs were Flag-tagged, anti-Flag antibody was used to examine CU-REWIREs and GAPDH was used as a loading control.

We then applied AlphaFold2 to predict structures of ProAPOBECs derived from human and rodent deaminase domains (Figures 2E and S2D). These ProAPOBECs showed structural conservation with human APOBEC3A, suggesting their potential functionality for base editing. We subsequently fused various ProAPOBECs-C171A with the ePUF10 domain, creating a new generation of CU-REWIREs (version 5.1 to 5.17) (Figures 2F, 2G and Table S1). Upon expression analysis, most of the CU- REWIRE5 variants (CU5s) demonstrated stable expression and various editing activity towards C_459_ site of EGFP mRNA expressed in the HEK293T cells (Figures 2H and 2I). This result demonstrated that we have significantly expanded the AID/APOBEC protein repertoire, leading to the development of novel RNA base editors with a broader targeting context for C-to-U editing. Next, we aim to characterize the editing properties and specificities of these newly engineered CU5s in additional target sites.

### Enhanced Targeting Range and Specificity in RNA Base Editing with CU5s Toolki

We evaluated the RNA editing activity of CU5s by testing their C-to-U editing capabilities at the artificially modified sites of EGFP mRNA, where the U_458_C_459_ was altered to GC, CC, and AC to test the sequence preference of CU5s (Figure 3A). Surprisingly, multiple CU5s base editor containing ProAPOBECs exhibited C-to-U transition under G<u>C</u>, C<u>C</u>, and A<u>C</u> contexts (Figure 3B). We found that while some new enzymes (like CU5.7 and 5.13) maintain the original preference of U<u>C</u> context for their editing activity (Figures 3B, S3A and S3B). The CU5.16, which demonstrates a particular interest in the novel C-to-U editors with A<u>C</u> context (Figures S3C-S3F).

**Figure 3.**
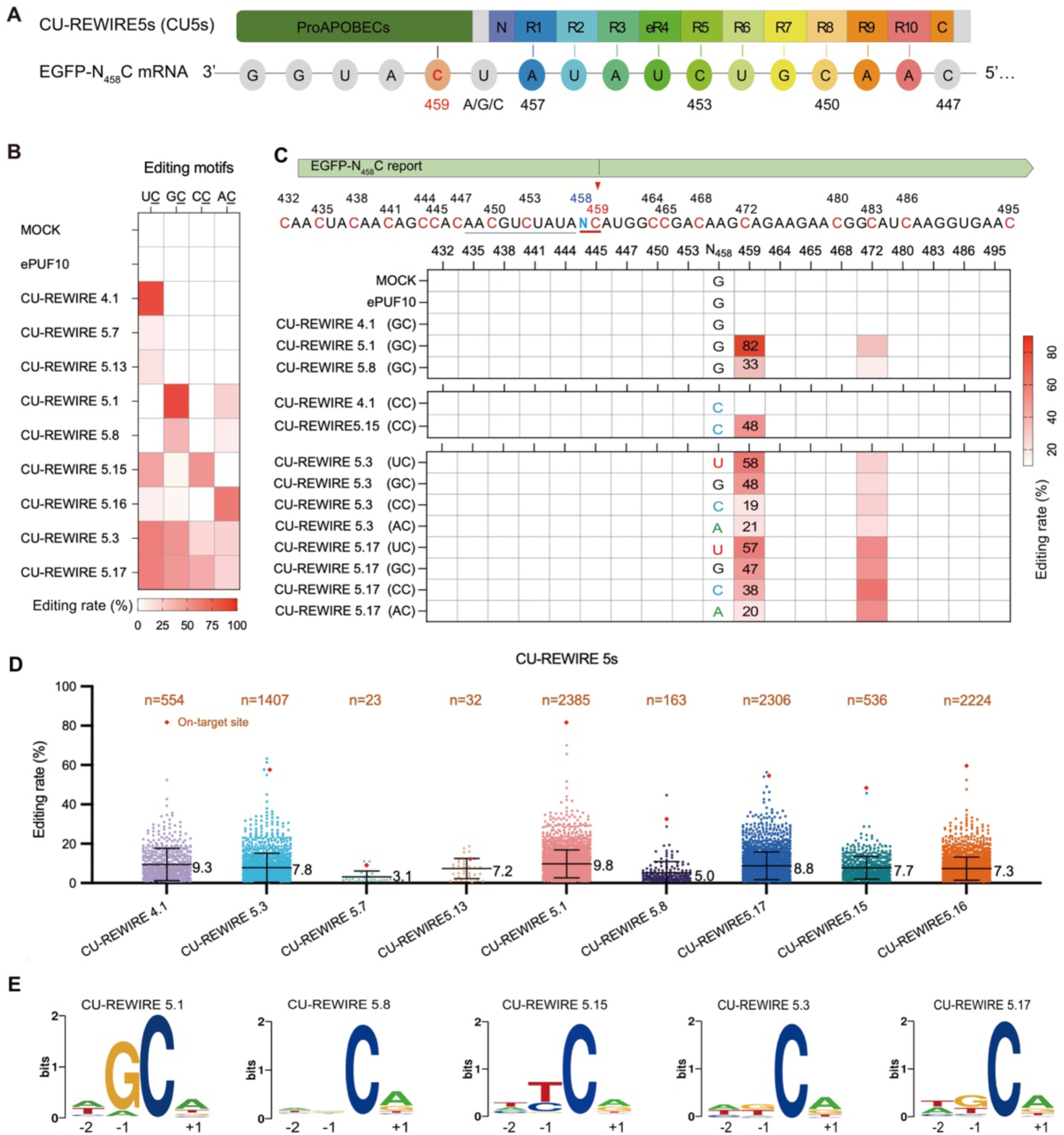
Enhanced targeting range and specificity in RNA base editing with CU5s. (A) Schematic illustration demonstrating the targeting of the C_459_ site in EGFP mRNA by CU5s. The native U_458_<u>C</u> context was manually altered to A_458_<u>C</u>, G_458_<u>C</u>, and C_458_<u>C</u> to test editing preferences. (B) Heatmap depicting the percent editing levels of CU5s at the C_459_ site of EGFP transcript in A<u>C</u>, U<u>C</u>, G<u>C</u>, and C<u>C</u> contexts. The PUF10 binding sequence and on-target site were showed by panel (A). (C) Editing efficacy of CU5s at the on-target and bystander site of EGFP transcript in A<u>C</u>, U<u>C</u>, G<u>C</u>, and C<u>C</u> contexts. A heatmap displays editing rates of all cytosines near the on-target site C_459_ for each CU5s and control. Editing rates of C_459_ are measured by RNA-seq with triplicates. (D) Global editing rates of transcriptome-wide RNA-seq by CU5s (related to data from panel B, with labels and cutoffs the same as in Figure 1G). Orange rhombuses indicate the on-target EGFP sites, and the average efficiency of total off-target editing events is noted on the right. n numbers represent edited cytosines detected. (E) Sequence motif logos derived from cytosines edited by different CU5s, based on RNA-seq data (related to panel D).

Importantly, new CU5s have high sequence specificity distinct from the original U<u>C</u> specificity ^21,27^. For example, the CU5.1 and 5.8, containing the deaminase domain of human APOBEC1 and APOBEC3H respectively, demonstrated significant C-to-U editing activity within G<u>C</u> or A<u>C</u> motifs (Figures 3B, S4A and S4B). Our finding underscores that Pro.hAPOBEC1 editing capabilities significantly differ from natural human APOBEC 3A (only editing U<u>C</u> motifs) and human APOBEC1 (editing U<u>C</u> motifs with low efficiency), indicating the emergence of novel cytidine editing activities in engineered ProAPOBECs. In addition, the CU5.15, containing the deaminase domain of mouse APOBEC3, demonstrated significant C-to-U editing activity within C<u>C</u> or U<u>C</u> motifs (Figures 3B and S4C). The other enzymes can edit cytosine at various sequence context, including two enzymes (CU5.3 and 5.17, containing the deaminase domain of mouse APOBEC1 and rat APOBEC1 respectively.) that can edit the cytosine in any context (i.e., N<u>C</u> motifs), suggesting they may function as a base editor with broad target range (Figures 3B, S4D and S4E). Such expansion of specificity is particularly useful in their application as a therapeutic protein (Figures 3B, S3 and S4).

We further used RNA-seq to assess the bystander editing effects of these new CU5s, with particular interest in the novel C-to-U editors with different sequence specificity (like GC editors CU5.1 and 5.8 (Figure 3C, top), the CC editor CU5.15 (Figure 3C, middle), and the NC editors CU5.3 and 5.17 (Figure 3C, below).

The RNA-seq experiments also allow us to assess the global off-target effects of CU5s across the entire transcriptome (Figure 3D). Our results demonstrated that the CU5.1, 5.17, and 5.16 showed higher off-targeting editing rates compared to CU4.1, while certain CU5s have lower number of off-target editing sites. There seems to be correlation between off-target editing rates and the on-target editing efficiency (i.e., more activity CU5s have a larger number of off-target edit sites), suggesting that we have to balance these two properties of CU5s in the practical applications. We further analyzed the off-target editing sites for their consensus sequence motifs (Figure 3E), and found that these motifs are largely consistent with the sequence preference observed using C-to-U editing on a single target site (C_459_) of EGFP mRNA (Figures 3B and S4).

### Editing Windows and *in vitro* Application of CU5s

To elucidate the editing characteristics of CU5s on the target RNA, we conducted a comprehensive evaluation of the editing windows associated with selected CU5s. Three variants, CU5.1, CU5.15 and CU5.17, were selected for this analysis, because they can edit cytosine in the new sequence context other that the original U<u>C</u> motifs. We found that CU5.1 (editing at G<u>C</u> motif) demonstrated the highest editing efficiency at positions 2- or 4-nucleotides (nt) downstream of the ePUF10 binding site in EGFP mRNA, as shown by different target reporters containing G<u>C</u> dinucleotides at different positions downstream of the ePUF10 binding site (Figure 4A).

**Figure 4.**
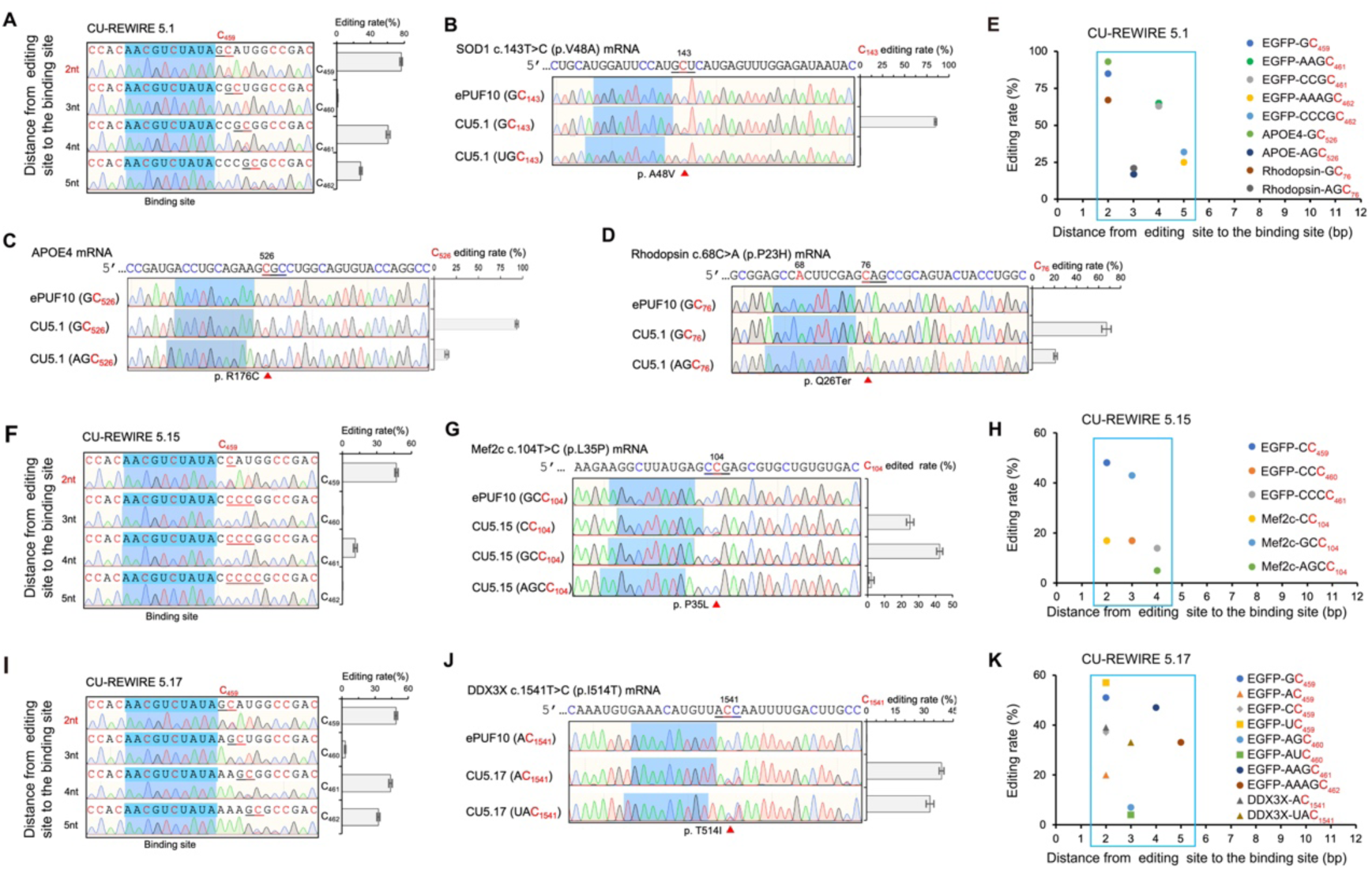
Editing windows and application of CU5s. (A) Determination of editing windows for CU5.1 on C_459_ of EGFP mRNA. The graph shows on-target editing rates by CU5.1 at various distances from the PUF binding site. (B-D), Editing efficacy of CU5.1 on C_143_ of SOD1 reporter mRNA (B), C_526_ of APOE4 mRNA (C) and C_76_ of Rhodopsin reporter mRNA (D) expressed in HEK293T cells. The PUF-binding sites are marked in blue. (E) Summary of the editing window for CU5.1 across EGFP, APOE4, and Rhodopsin reporter mRNAs. The x-axis represents the distance from the editing site to the PUF binding site; the y-axis shows the C-to-U editing rate. Editing rates were acquired by Sanger sequencing. (F) Determination of editing windows for CU5.15 on C_459_ of EGFP mRNA. (G) Editing efficacy of CU5.15 on C_104_ of Mef2c reporter mRNA expressed in HEK293T cells. (H) Summary of the cytosine editing window mediated by CU5.15 around the target sites of EGFP and Mef2c reporter mRNAs. (I) Determination of editing windows for CU5.17 on C_459_ of EGFP mRNA. (J) Editing efficacy of CU5.17 on C_1541_ of DDX3X reporter mRNA expressed in HEK293T cells. (K) Summary of the cytosine editing window mediated by CU5.17 around the target sites of EGFP and DDX3X reporter mRNAs. Values represent mean ± SEM (n=3).

We further evaluated the efficacy of CU5.1 in reporter genes carrying disease-causing mutations. For instance, the T143C mutation in the *SOD1* allele, which results in the p.V48A substitution in the SOD1 protein, is the most common mutation associated with ALS in Southeastern China ^44^. We designed two CU5.1 enzymes with ePUF10 domains targeting sequences at positions U_132_–U_141_ and A_131_–A_140_ (i.e., 2 or 3 nucleotides upstream of C_143_, respectively). The CU5.1 enzyme binding 2 nucleotides upstream of the mutated site successfully converted C_143_ to U_143_ with an editing efficiency of ∼85%, effectively transforming the mutant *SOD1* allele into the benign form ^45^ (Figure 4B).

In another example, the T526C mutation in the APOE4 allele, causing p.R176C in APOE4 protein, was found to be associated with Alzheimer’s disease ^46,47^. We designed CU5.1 binding to the APOE4 sequences, and found that the CU5.1 binding at 2-nt upstream of the mutated site effectively converted C_526_ to U_526_ with editing efficiency at ∼90%, potentially transforming APOE4 into the benign APOE1 allele ^48^ (Figure 4C). In addition, the c.68C>A (p.P23H) mutation in the human Rhodopsin gene, known to cause Retinitis Pigmentosa (RP) ^49^ can be effectively edited by CU5.1 that bind to 2-nt upstream of the mutated site (Figure 4D). As a result of editing C_76_ to U_76_, we introduced a stop codon in the mutated mRNA, which should abolish the toxic protein mutant (Figure 4D). Combining the results across different target mRNAs, we summarized the editing window for CU5.1 as 2∼5-nt downstream of the ePUF10 binding site (Figure 4E).

Furthermore, CU5.15 demonstrated the highest editing efficiency 2-nt downstream of the ePUF10 binding site in EGFP mRNA (Figure 4F). We adopted similar approach to examine the editing window of CU5.15 using reporters containing C<u>C</u> at different distance from the designed binding site (Figure 4F), or using multiple CU5.15 targeting the c.104T>C (p.L35P) mutation in MEF2C gene (Figure 4G) ^8^. We found that CU5.15 effectively edited cytidine 2-nt or 3-nt downstream of the ePUF10 binding site, resulting in an editing window within 2∼4-nt in various target mRNAs (Figure 4H).

Finally, CU5.17, with editing activity in “N<u>C</u>” context, demonstrated a high efficiency at positions 2-nt downstream of the ePUF10 binding site as judged by EGFP reporter (Figure 4I). Importantly, CU5.17 was highly efficient in editing the T1541C site of the human DDX3X mRNA linked to neurodevelopmental disorders ^50^, particularly when the CU5.17 binding to 2-nt or 3-nt upstream of the mutated site (Figure 4J). Although bystander editing at C_1542_ occurred, this did not affect the amino acid sequence of DDX3X (Figure 4J), indicating the editing window for CU5.17 is within 2∼5-nt after the ePUF10 binding site across different mRNAs (Figure 4K).

### Improving the CU5 system with AI-Assisted Design of ProAPOBECs utilizing cytosine Deaminase derived from Mammals

We also explored the possibility of using ProAPOBECs derived from deaminase of other mammalian species using similar approach (Figures S5 and Table S1). AlphaFold2-based predictions revealed that the deaminase domains from many mammalian APOBEC1s and APOBEC3s may form functional structures with the NTD and CTD of the human APOBEC3A (Figure 5A). We found that both Z1 and Z2 domains of mammalian APOBEC3s could form functional ProAPOBECs when combined with NTD and CTD of human APOBEC3A (Figures S5B and S5C).

**Figure 5.**
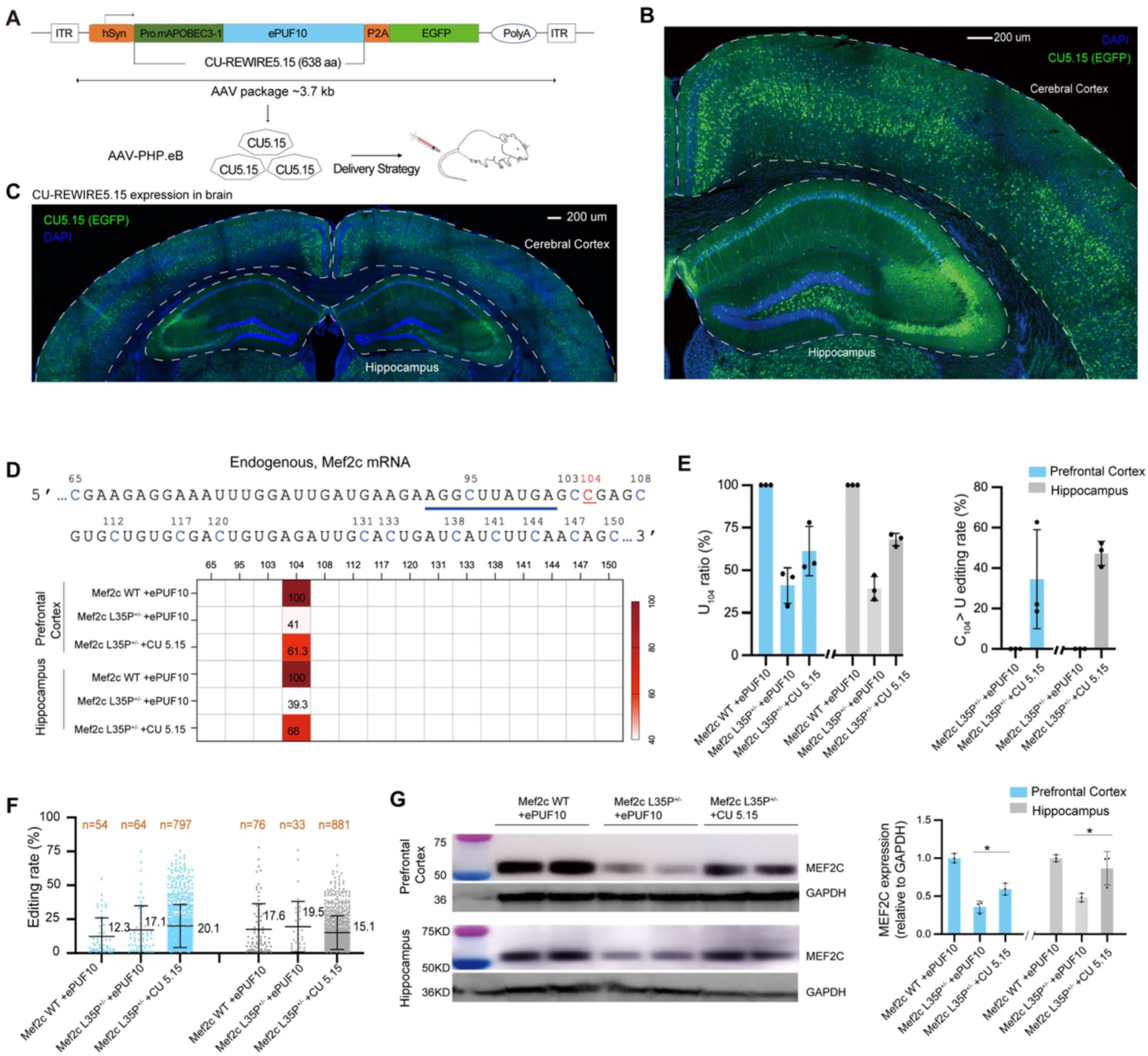
*In vivo* RNA base editing in Mef2c L35P^+/−^ mice corrects genetic mutations and protein expression. (A) Schematic illustration of AAV-CU-REWIRE5.15 (CU5.15) vectors delivered into Mef2c L35P^+/−^ mice through intravenous injection of AAV-PHP.eB virus in the tail vein. (B) Immunofluorescence images showing CU5.15 (green) and DAPI (blue) in the cortical and hippocampal brain regions, 4-weeks post-viral injection. Scale bar: 200µm. (C) Immunofluorescence images of CU5.15 (green) and DAPI (blue) in the cortical and hippocampal regions, 16-weeks post-viral injection. (D) Editing efficiency of C_104_ in endogenous mouse Mef2c mRNA by CU5.15. The ePUF10 binding site is underlined in blue, C_104_ marked in red (top). Lower panel: Editing rates of cytosines around the on-target site. Rates were measured by RNA-seq with triplicates. Values represent mean ± s.e.m. (n = 3). (E) Percentage of U_104_ in prefrontal cortex and hippocampus tissues of Mef2c WT or L35P^+/−^ mice edited by ePUF10 or CU5.15, measured by RNA-seq with triplicates (Left). Editing efficiency of C-to-U of the Mef2c C_104_ site by CU5.15 in various groups (Right). Editing rate formula: C_edited_ =(U_CU5.15_-U_ePUF10_) / C_total_ (1-U_ePUF10_). (F) Scatter plots showing transcriptome-wide C-to-U RNA base editing in prefrontal cortex or hippocampus samples treated with CU5.15 and ePUF10 control in mice brain samples. Values represent mean ± s.e.m. (n = 3). (G) MEF2C protein levels in the prefrontal cortex or hippocampus of Mef2c WT or L35P^+/−^ mice treated with ePUF10 or CU5.15. anti-MEF2C antibody was used. GAPDH used as a loading control (Left). Quantification of MEF2C protein level (Right). ePUF10 vs. CU5.15 in prefrontal cortex, P = 0.0203; ePUF10 vs. CU5.15 in hippocampus, P = 0.0423), Unpaired two-sided Student’s t-test was used, statistical values represent the mean ± s.e.m. *P < 0.05. n = 3 mice.

Furthermore, we integrated these newly engineered ProAPOBECs into CU-REWIRE (i.e., by fusing with ePUF10 domain) and tested their editing activity on EGFP mRNA. We found significant C-to-U editing activity in G<u>C</u>, A<u>C</u>, and C<u>C</u> contexts by different ProAPOBECs (Figures S5D and S5E). ProAPOBECs with deaminase domains from APOBEC1 showed strong activity in G<u>C</u> contexts, while some with APOBEC3-Z1 (A3Z1) domains demonstrated activity in C<u>C</u> and A<u>C</u> contexts (Figures S5F-S5H). These results highlight the versatility of engineered ProAPOBECs for C-to-U base editing, suggesting an expansion of specificity is particularly useful in potential therapeutic applications.

### *In Vivo* RNA Editing in an ASD Mouse Model Using CU5.15

Effective RNA base editing in the brain has been a challenging endeavor ^20^. To explore the potential of CU5s, powered by ProAPOBECs, for *in vivo* RNA editing, we utilized an ASD mouse model with L35P (c.104T>C) point mutation in MEF2C gene, which was reported to cause severe ASD in human ^8^. An APOBEC-embedded cytosine base editor for DNA has successfully performed C-to-T conversion at the C_104_ site of the Mef2c gene in the Mef2c L35P mutant mice, effectively reversing the ASD-like phenotypes in this mouse model ^8^. However, the previous methods need to co-inject two AAV vectors for in vivo DNA base editing, posing a practical challenges for its application in clinical settings ^6,8,51^. Packaging the base editing system into a single AAV vector could significantly reduce costs and enhance delivery efficiency in clinical applications.

We first examined the C-to-U base editing efficiency of the CU-REWIRE system as compared with Cas13-based RNA editing tools on the C_104_ site on the Mouse Mef2c (c.104) report mRNA ^23,27^ (Figure S6). We observed that CU5s exhibited significantly higher editing efficiency compared to Cas13-based tools. Specifically, CU5.15 achieved an editing efficiency of 43%, and CU1.15 reached 18% (Figures S6A and S6B), whereas the Cas13-based tool CURE-X-Pro.mAPOBEC3-1 showed only 5% efficiency (Figures S6C and S6D). Notably, the CURE-X system utilizing native APOBEC proteins failed to edit the target *Mef2c* mRNA. However, replacing the native APOBEC with ProAPOBEC in the CURE-X-Pro.mAPOBEC3-1 system resulted in effective editing (Figures S6C-S6E), underscoring the therapeutic potential of ProAPOBECs in Cas13-based RNA base editors.

We further packaged CU5.15, featuring the ePUF10 module targeting the A_92_-A_101_ region of mouse Mef2c mRNA, into brain blood barrier (BBB)-crossing AAV-PHP.eB vectors (Figure 5A). These vectors, along with an EGFP reporter driven by the human synapsin 1 gene promoter (hSyn), were injected into the tail vein of 4-week-old Mef2c WT and L35P^+/−^ mice ^8^. Robust EGFP expression was observed in the cortical and hippocampal regions 4- and 16- weeks post-injection (Figures 5B and 6C), confirming the efficient gene delivery in brains.

**Figure 6.**
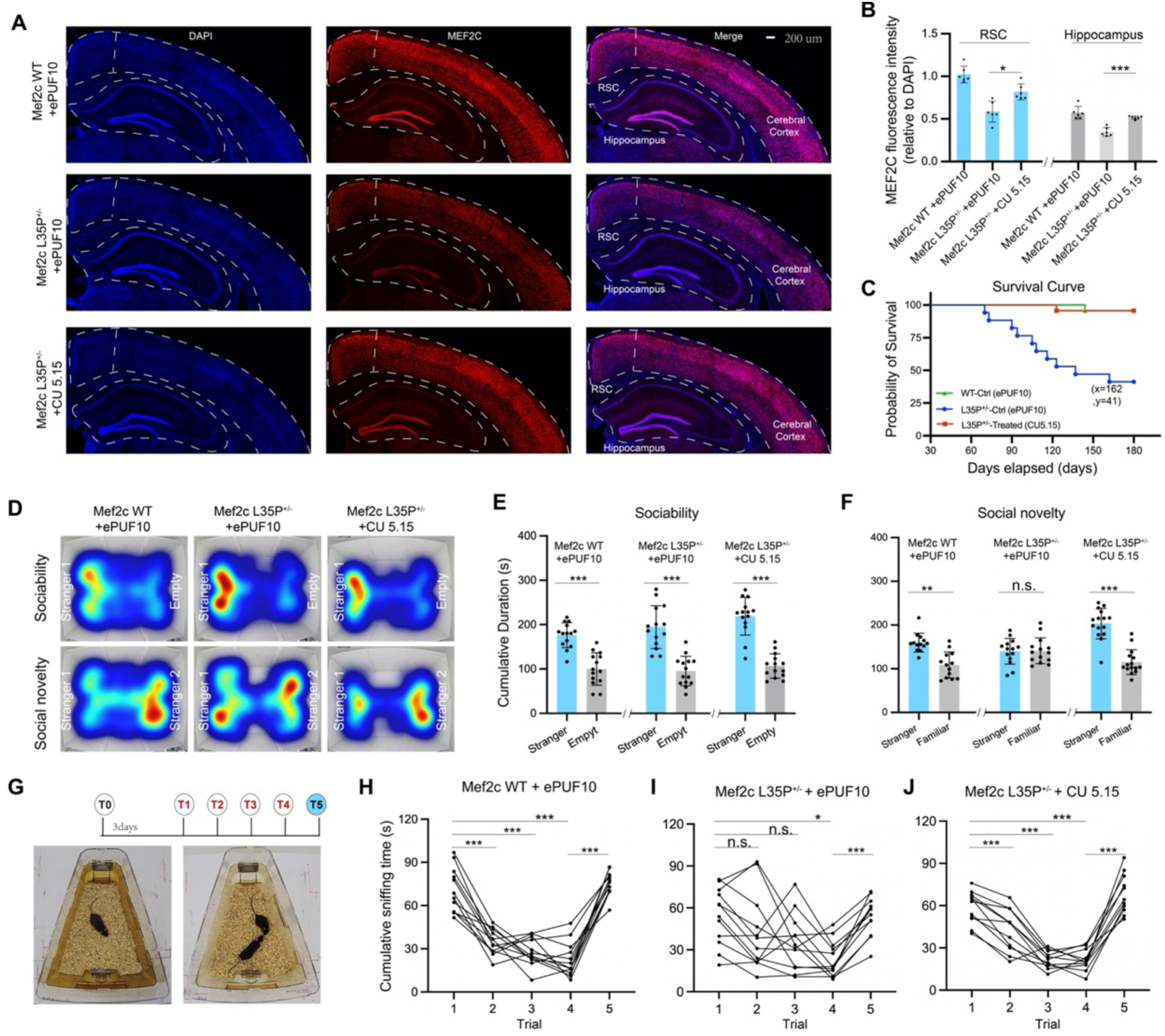
*In vivo* RNA base editing in Mef2c L35P^+/−^ mice rescued abnormal social behaviors. (A) Immunohistochemical staining for MEF2C (red) and DAPI (blue) in the retro splenial cortex (RSC) and hippocampus of Mef2c WT and L35P^+/−^ mice, indicating viral injections. (B) Left: Quantification of MEF2C protein fluorescence density in the RSC (ePUF10 vs. CU5.15, P = 0.0040). Right: In the hippocampus (ePUF10 vs. CU5.15, P < 0.0001. *P < 0.01. n = 6 brain slices from three mice of each group). (C) Lifespan of Mef2c L35P^+/−^ mice with ePUF10 or CU5.15. Kaplan-Meier curve statistical analysis (L35P^+/−^(ePUF10) versus L35P^+/−^(CU5.15), Chi square=25.99, P < 0.0001). Kaplan-Meier curve statistical analysis, Log-rank (Mantel-Cox) test was used. WT (ePUF10) n = 23 mice, L35P^+/−^ (ePUF10) n = 17 mice, L35P^+/−^ (CU5.15) n = 23 mice). (D) Heatmaps representing locomotion of Mef2c WT or L35P^+/−^ mice in the sociability and social novelty sessions of the three-chamber test (E) Cumulative interaction duration with mouse and empty cage in different groups of Mef2c mice (stranger vs. empty, WT+ePUF10, P < 0.0001, L35P^+/−^+ePUF10, P < 0.0001; L35P^+/−^+ CU5.15, P < 0.0001. n = 14 mice). (F) Cumulative interaction duration with stranger and familiar mouse in different groups of Mef2c mice (stranger vs. familiar, WT+ePUF10, P = 0.0007; L35P^+/−^+ePUF10, P = 0.9130; L35P^+/−^+ CU5.15, P < 0.0001. Significantly different, P < 0.01. n = 14 mice). (G) Diagram of the home cage experiment. T0: social isolation for three days. T1-T4: Social interaction trials with one same partner mouse. T5: Social interaction with a novel partner mouse. Recording sniffing time within a 2-minute period. (H) Sniffing time in social intruder test for Mef2c WT +ePUF10 (T1 vs. T2, P < 0.0001; T1 vs. T3, P < 0.0001; T1 vs. T4, P < 0.0001; T4 vs. T5, P < 0.0001. n = 12 mice). (I) Sniffing time in the social intruder test for Mef2c L35P^+/−^ +ePUF10 (T1 vs. T2, P = 0.0153; T1 vs. T3, P = 0.0614; T1 vs. T4, P = 0.0026; T4 vs. T5, P < 0.0001. n = 12 mice). (J) Sniffing time in the social intruder test for Mef2c L35P^+/−^ +CU5.15 (T1 vs. T2, P < 0.0001; T1 vs. T3, P < 0.0001; T1 vs. T4, P < 0.0001; T4 vs. T5, P < 0.0001. n = 12 mice). Paired two-sided Student’s t-test was used, statistical values represent the mean ± s.e.m. * P < 0.01, ** P < 0.001, *** P < 0.0001.

Sanger sequencing and RNA-seq of prefrontal cortical and hippocampus tissue samples from the mice revealed that CU5.15 induced C-to-U editing at the C_104_ site in the Mef2c L35P^+/−^ mice injected with CU5.15 (Figure S7). RNA-seq experiments showed a significant increase in the percentage of U_104_ in the prefrontal cortex (61.3%) and hippocampus (68%) of Mef2c L35P^+/−^ mice injected with CU5.15, compared to those injected with ePUF10 (Figures 5D and 6E). This data confirmed the induction of C-to-U editing events by CU5.15. Additionally, no bystander editing events were observed in the C nucleotides from positions C_65_-C_150_ of Mef2c mRNA, demonstrating the precision of CU5.15-mediated RNA editing (Figure 5D). C-to-U editing rates across various brain regions ranged from 30-50% (Figure 5E). Considering that the brain tissues collected for RNA-seq are mixture of neurons and glial cells, the exact C-to-U editing of the Mef2c L35P site in neurons would be higher. Transcriptome-wide RNA sequencing revealed approximately 800 C-to-U editing events associated with CU5.15 (Figure 5F).

Notably, MEF2C protein levels were fully restored in the cortical and hippocampal regions of Mef2c L35P^+/−^ mice injected with CU5.15, as compared to those injected with ePUF10 and to Mef2c WT mice (Figure 5G). This restoration was further validated by immunostaining using an anti-MEF2C antibody (Figures 6A and 6B). Remarkably, the lifespans of Mef2c L35P^+/−^ mice injected with CU5.15 were close to WT group (Figure 6C), which was not observed in previous DNA editing work ^8^, suggesting that RNA base editing facilitated by CU5.15 may exert a broader impact in this mouse model.

We subsequently examined the influence of CU5.15 on autistic-like behaviors in Mef2c L35P mice using a series of social interaction tests. The investigation began with the three-chamber test, comprising both sociability and social novelty tests. In the sociability test, mice were given the choice between interacting with another mouse or an empty cage (Figure 6D). This was followed by the social novelty test, where the mice had to choose between a familiar partner and a novel partner. Mice treated with AAV-CU5.15 demonstrated a significant improvement in social interaction performance during the social novelty test ^8^ (Figures 6E and 6F).

The study progressed to analyzing social interaction behaviors using the social intruder test, consisting of six trials (T0-T5) (Figure 6G). After three days of social isolation (T0), the mice encountered the same partner for four consecutive trials (T1-T4, each lasting 5 minutes), and then a new partner in the fifth trial (T5). The duration of sniffing time between the subject mice and their partners was recorded. Notably, the aberrant social behaviors observed in Mef2c L35P mice during the initial four trials (T1-T4) with intruder partners were entirely mitigated following treatment with CU5.15 (Figures 6H-6J). These findings suggest that in vivo RNA base editing facilitated by CU5.15 effectively restores normal MEF2C protein expression and alleviates the social impairments associated with the Mef2c L35P mutation.

In summary, our study demonstrates that RNA base editing, as facilitated by CU5.15, provides a potent tool for correcting genetic mutations in the brain and has potential therapeutic applications for ASD and other genetic disorders.

## Discussion

APOBEC3A is a highly active deaminase frequently used in DNA base editors. However, its application has been hindered by some intrinsic limitations of this enzyme, including off-target effects and RNA binding issues ^11,12^. Although point mutations in APOBEC3A have reduced off-target effects in DNA base editing, its application in RNA base editing has remained largely unexplored ^52,53^. We hypothesized that the high off-targeting rates of APOBEC3A might be largely due to its close interaction with target mRNA. Structurally, the dimerization of APOBEC3A plays a key role in stabilizing its mRNA binding. Therefore, our initial strategy was to modulate dimerization to minimize its off-target effects and decrease the strong associations with mRNA. Indeed, APOBEC3A mutants in the NTD or CTD, exhibiting reduced mRNA interaction and dimerization, showed fewer off-target effects. This provided a range of APOBEC3A variants for precise RNA base editing in conjunction with the REWIRE system (Figures 1C-1I). However, such variants were limited to editing cytidines in UC contexts (Figure S2A), indicating the need for a more fundamental protein engineering of cytosine base editors with broader targeting scopes.

Utilizing AlphaFold2-assisted protein structure analysis, we discovered that the deaminase domains of APOBECs could act as interchangeable modules (Figure 2). This suggested that the NTD and CTD may contribute to the recognition of viral genetic materials in various organisms. Consequently, we anticipated that novel deaminase activities could emerge from new combinations of deaminases with NTDs and CTDs. This led to the identification of a series of APOBECs with novel cytidine-editing activity under various contexts, significantly broadening the application of REWIRE- mediated RNA base editing.

Compared to Cas13-based C-to-U base editors, the CU5s with installed ProAPOBECs demonstrated an impressive capacity for effecting C-to-U transitions. Notably, our study revealed that the CU 5.7 and 5.13 are capable of editing cytosine in U<u>C</u> motifs, akin to the CasR13-based RNA editing system that utilizes human APOBEC3A or evolved ADAR2 enzyme ^21,23^, the CU5.15 enzyme can edit cytosine across C<u>C</u> sequence context, similar to the SNAP-CDAR-S system that employs a evolved ADAR2 enzyme ^24^. Meanwhile, the CU5.3 and 5.17 enzyme can edit cytosine across N<u>C</u> sequence context, suggesting their potential as broad-spectrum editing tools. Importantly, multiple CU5s base editor have shown remarkable C-to-U transitions capabilities within G<u>C</u>, A<u>C</u>, and C<u>C</u> motifs, exhibiting high sequence specificity distinct from traditional RNA base editors (Figure 3). For example, CU5.1 has shown significant C-to-U editing activity within G<u>C</u> motifs, albeit with lower efficiency in A<u>C</u> motifs. This enzyme notably reduces bystander editing effects and enhances editing specificity in the target region. Such expansion of specificity is particularly useful in their application as a therapeutic protein (Figures 3, S3 and S4).

While Cas13-mediated A-to-I RNA editing has been reported in mouse models of human genetic disorders ^18,20^, effective C- to-U RNA editing had not been achieved due to the intrinsic properties of APOBECs. With the new REWIRE system, we demonstrated that ProAPOBEC can effectively perform RNA base editing in the liver and brain (Figures 5-6). With an editing rate of nearly 50% *in vivo*, Mef2c mutant mice showed restored protein levels and correction of autistic-like behaviors. Furthermore, effective RNA base editing prolonged the lifespan of Mef2c mutant mice, a result not observed with the Mef2c L35P DNA editing ^8^. The effects of *in vivo* RNA base editing in our study appear to be long-lasting for over 6 months (Figure 6C).

ProAPOBECs also have potential applications in the field of RNA epigenetics. The elucidation of epigenetic modifications in DNA and RNA has become crucial for understanding their diverse biological functions ^54^. RNA modifications add a new layer to gene regulation, giving rise to “RNA epigenetics.” Techniques such as DART-seq (deamination adjacent to RNA modification targets) ^55^, ACE-Seq (APOBEC-coupled epigenetic sequencing) ^56^, and AMD-Seq (APOBEC3A-mediated deamination sequencing) ^57^, all employing APOBECs manipulation, have been developed for comprehensive mapping of RNA epigenetics. Additionally, RNA-binding proteins (RBPs) are pivotal in gene expression and RNA processing. To elucidate RBP dynamics at the single-cell level, the Gene lab developed STAMP (Surveying Targets by APOBEC-Mediated Profiling), enhancing the detection of RBP-RNA interactions ^58^.

The ProAPOBECs engineered in this study have enhanced editing efficiency and specificity over their natural APOBEC counterparts, emerging as significant improvement to the current repertoire of tools available for genetic modification. Our objective is to refine the ProAPOBECs’ activity and expand their application, facilitating the development of streamlined and accurate methods for conducting both DNA and RNA edits in vivo. In summary, the integration of REWIRE technology with ProAPOBEC-mediated base editing heralds a promising new era in both fundamental research and applied therapeutic strategies. This work not only enriches the existing collection of tools for genetic engineering but also paves the way for novel approaches to treat and manage genetic disorders, marking a significant step forward in the field of genomics and molecular medicine.

## Acknowledgments

We extend our gratitude to Dr. Li Yang of Fudan University for the gift of native human/mouse APOBECs, to Dr. Tian Chi of ShanghaiTech University for the CURE plasmids, and to Dr. Chunlong Xu of Lingang Laboratory for the xCBE plasmids. This work is supported by Fundamental Research Funds for the Central Universities (Z.Q), NSFC grants (82430046 to Z.Q., 32030064 to Z.W., 82201632 to W.H.); the Science and Technology Commission of Shanghai Municipality (2018SHZDZX05), Innovative research team of high-level local universities in Shanghai (SHSMU-ZDCX20211100) (Z.Q); the National Key Research and Development Program of China (2021YFA1300503), the Starry Night Science Fund at Shanghai Institute for Advanced Study of Zhejiang University (SN-ZJU-SIAS-009) (Z.W.). W.H. is supported by the Fellowship of China Postdoctoral Science Foundation (2022M723244) and the Shanghai Postdoctoral Excellence Program (2022777).

## Author contributions

Conceptualization, W.H., Z.Q., Z.W.; Biochemical and cell culture experiments, W.H.; NGS data analysis, B.Y.; AI-Assisted protein structure predictions, X.F., M.H.; Animal experiments, W.H., W.L., Y.Y., Y.Z., S.W. and S.S.; Writing, W.H., Z.W. and Z.Q.

## Data and code availability

The original RNA sequencing datasets have been deposited in the NCBI Gene Expression Omnibus (accession code PRJNA1068267). Custom computer code is available upon request. Accession numbers of RNA-seq data are provided in Table S2.

