## supplementary files for "Effective *in vivo* RNA base editing *via* engineered cytidine deaminase APOBECs fused with PUF proteins"

#### This Supplementary file includes:

**Supplementary Figures S1 to S7**

**Supplementary Tables S1and S2**

### Supplementary Figures and Legends

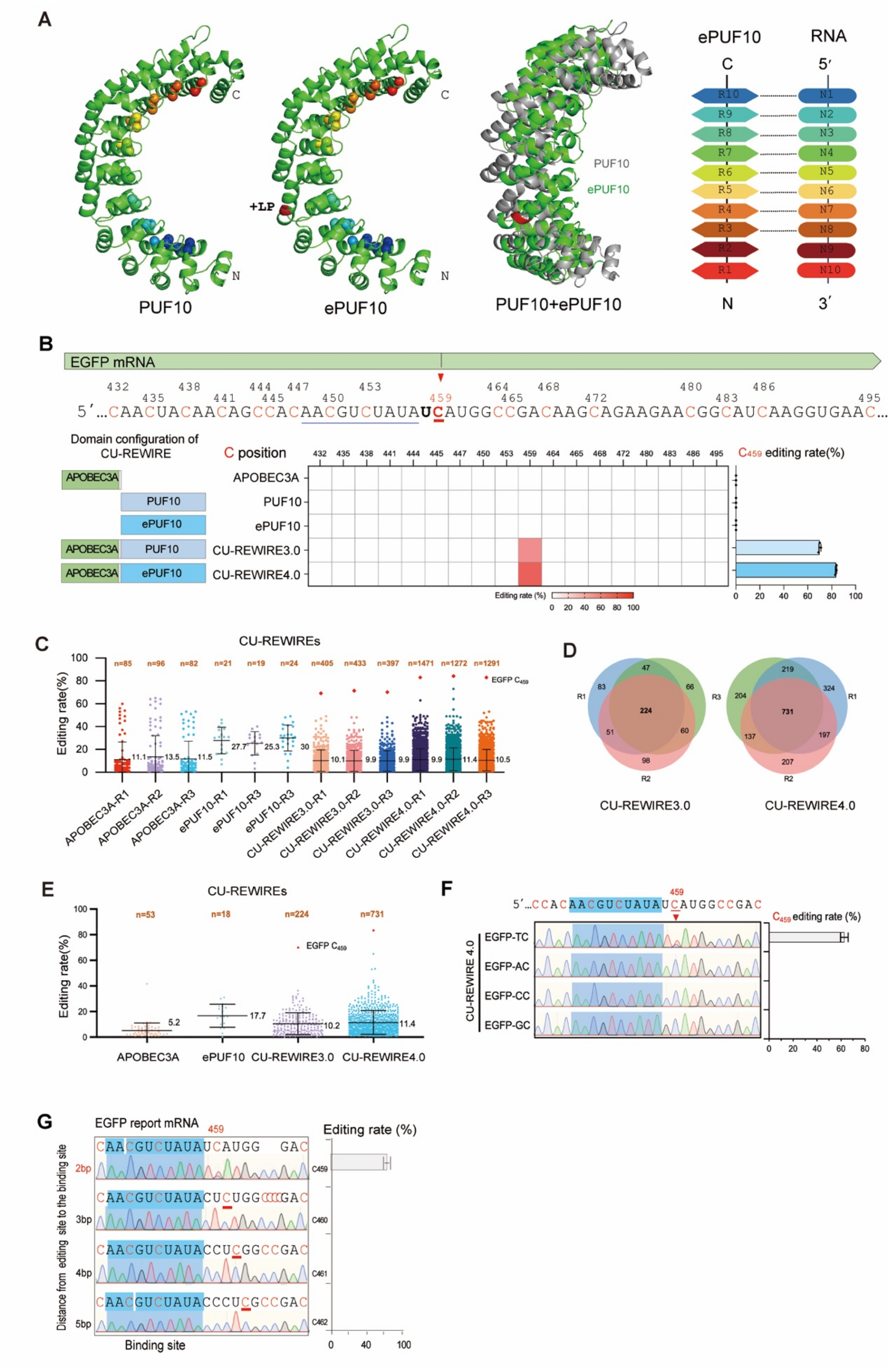

**Figure S1. Structure-based optimization of the PUF10 domain enhances stability and efficacy of CU-REWIRE.**

(A) AlphaFold2-predicted structure of the PUF10 domain (left) and a diagram illustrating PUF10-RNA interaction (right). ePUF10 recognizes individual RNA nucleosides through the Watson-Crick edge.

(B) Efficient editing of $C_{459}$ in EGFP mRNA by CU-REWIRE3 and CU-REWIRE4. The PUF10 binding site in EGFP is underlined in blue, with adjacent cytosines marked in red (top). A heatmap displays editing rates of all cytosines near the on-target site $C_{459}$ for each CU-REWIRE and control. Editing rates of $C_{459}$ are shown on the right, measured by RNA-seq with triplicates (see methods).

(C) Global editing efficiency of C-to-U RNA editing in samples treated with CU-REWIRE variants (related to data from panel B). Orange rhombuses indicate the on-target site EGFP-$C_{459}$, and the average efficiency of total off-target editing events is noted on the right. 'R' denotes replication, and 'n' represents the total number of edited cytosines detected.

(D) Venn diagram illustrating the number of "C-to-U changes" (editing noise, see methods) in different versions of CU-REWIREs (data from panel C).

(E) Transcriptome-wide editing events of C-to-U base editing in samples treated with CU-REWIRE3 and 4 variants (data from D).

(F) Editing activity of CU-REWIRE4.0 on the $C_{459}$ site of the EGFP transcript, considering different adjacent bases.

(G) On-target editing rates of CU-REWIRE4.0 at various distances from binding sites, representing the number of bases between the editing site and the 3′ end of the PUF binding site. Values represent mean ± SEM (n=3).

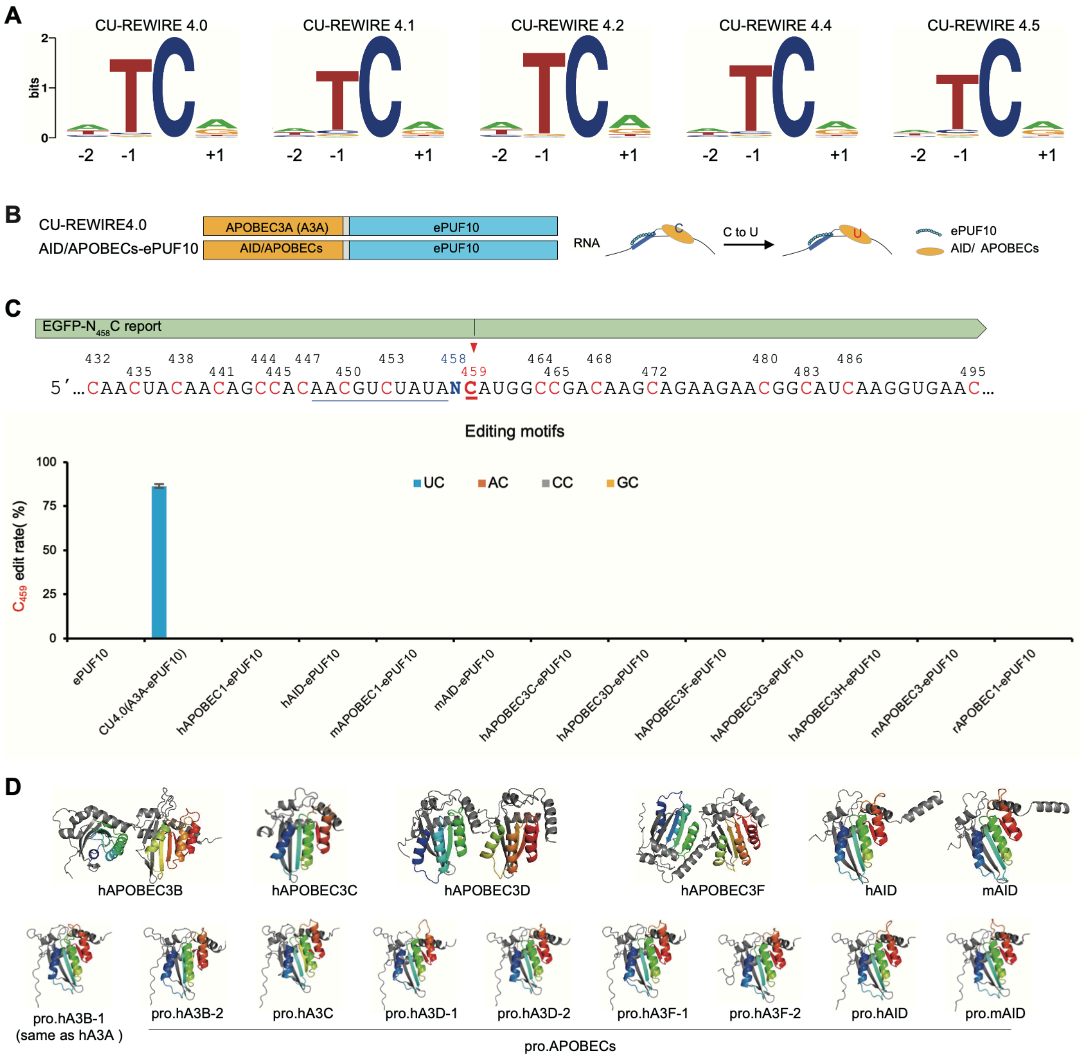


**Figure S2. Systematically examining various AID/APOBEC enzymes that are integrated within the CU-REWIRE platform.**

(A) Sequence logos derived from edited cytosines, based on RNA-seq data from samples treated with different CU-REWIRE variants (data from Figure 1H).

(B) Engineering of AID/APOBECs-ePUF10 editors for C-to-U editing. Left: Domain configuration of AID/APOBECs-ePUF10 with various native AID/APOBEC protein versions; Right: Schematic illustrating AID/APOBECs-ePUF10 mediated specific C-to-U base editing on RNAs.

(C) Top, AID/APOBECs-ePUF10 editors targeting the $C_{459}$ site of EGFP mRNA expressed in HEK293T cells. To evaluate editing efficiency, $N_{458}$ point mutations were introduced in EGFP, generating variants EGFP-$A_{458}$C, EGFP-$C_{458}$C, EGFP-$G_{458}$C, and EGFP-$U_{458}$C, each altering the context of the on-target site $C_{459}$. The PUF binding site in EGFP-$N_{458}$C reporter and adjacent cytidines are marked. Bottom, efficient editing of EGFP-$N_{458}$C mRNA by various AID/APOBECs-ePUF10 versions. For each version, the editing rate of $C_{459}$ was measured by Sanger sequencing, with CU-REWIRE4.0 (CU4.0) serving as a positive control. Values represent mean ± SEM (n=3).

(D) Alphafold2-assisted design of AID/APOBECs. Top, AlphaFold2-predicted structures of native AID/APOBECs, bottom, AlphaFold2-predicted structures of ProAPOBECs.

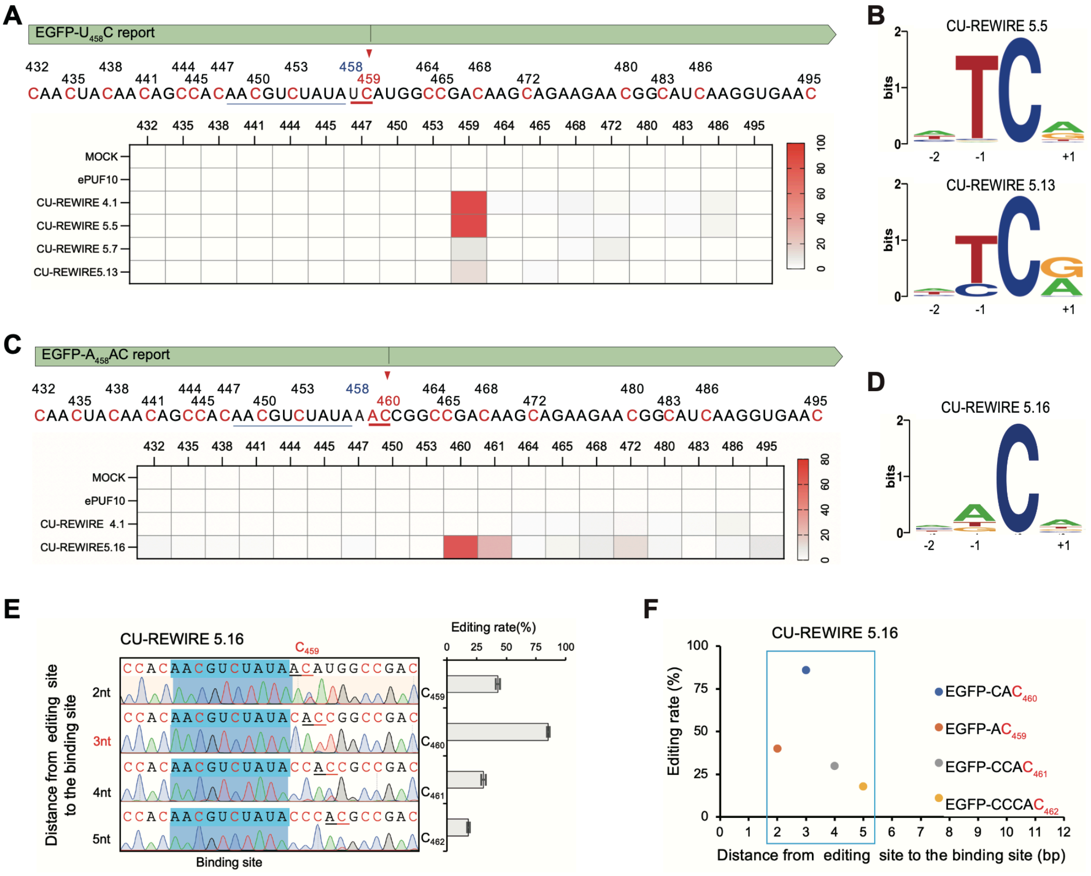


**Figure S3. Editing preferences and windows of CU-REWIRE5 variants.**

(A) Editing preferences of EGFP $C_{459}$ in the UC context by CU-REWIRE 4.1, CU-REWIRE 5.5 (similar as CU-REWIRE4.1, see Supplementary Figure 2D), 5.7 and 5.13. The ePUF10 binding site in EGFP is underlined in blue (top). A heatmap displays editing rates of all cytosines near the on-target site $C_{459}$ for each CU-REWIRE and control. Editing rates of $C_{459}$ are measured by RNA-seq with triplicates.

(B) Sequence motif logos derived from cytosines edited by CU-REWIRE5.5 and 5.13, based on RNA-seq data (related to Figure 3D).

(C) Editing preferences of EGFP $C_{460}$ in the AC context by CU-REWIRE5.16 and 4.1. Editing rates of $C_{460}$ are measured by RNA-seq with triplicates.

(D) Sequence motif logos derived from cytosines edited by CU-REWIRE5.16, based on RNA-seq data (related to Figure 3D).

(E) Determination of editing windows for CU-REWIRE5.16 in EGFP. The graph displays on-target editing rates by CU-REWIRE5.16 at various distances from the PUF binding site.

(F) Summary of the editing window for CU-REWIRE5.16 in EGFP. The x-axis represents the distance from the editing site to the PUF binding site, and the y-axis shows the C-to-U editing rate.

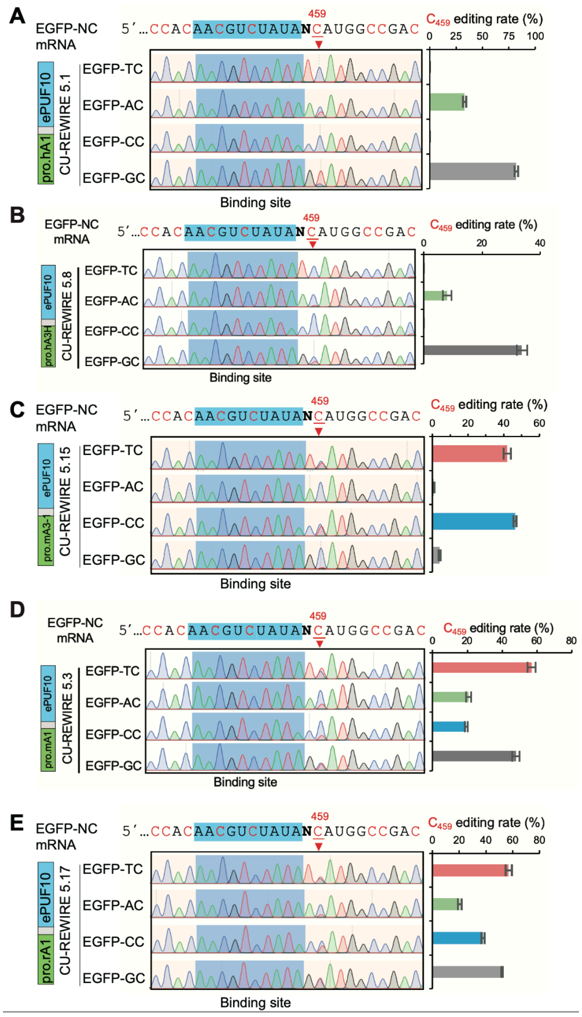


**Figure S4. Editing preferences of CU-REWIRE5 variants.**

(A) Editing efficacy of CU-REWIRE5.1 at the $C_{459}$ site of EGFP mRNA in $U_{458}C$, $A_{458}C$, $C_{458}C$, and $G_{458}C$ contexts. PUF binding sites in EGFP mRNA are highlighted in blue. Base editing rates were measured by Sanger sequencing.

(B) Editing efficacy of CU-REWIRE5.8 at the $C_{459}$ site in EGFP mRNA across UC, AC, CC, and GC contexts.

(C) Editing efficacy of CU-REWIRE5.15 at the $C_{459}$ site in EGFP mRNA in UC, AC, CC, and GC contexts.

(D), Editing efficacy of CU-REWIRE5.17 at the $C_{459}$ site in EGFP mRNA across UC, AC, CC, and GC contexts.

(E) Editing efficacy of CU-REWIRE5.3 at the $C_{459}$ site in EGFP mRNA in UC, AC, CC, and GC contexts.

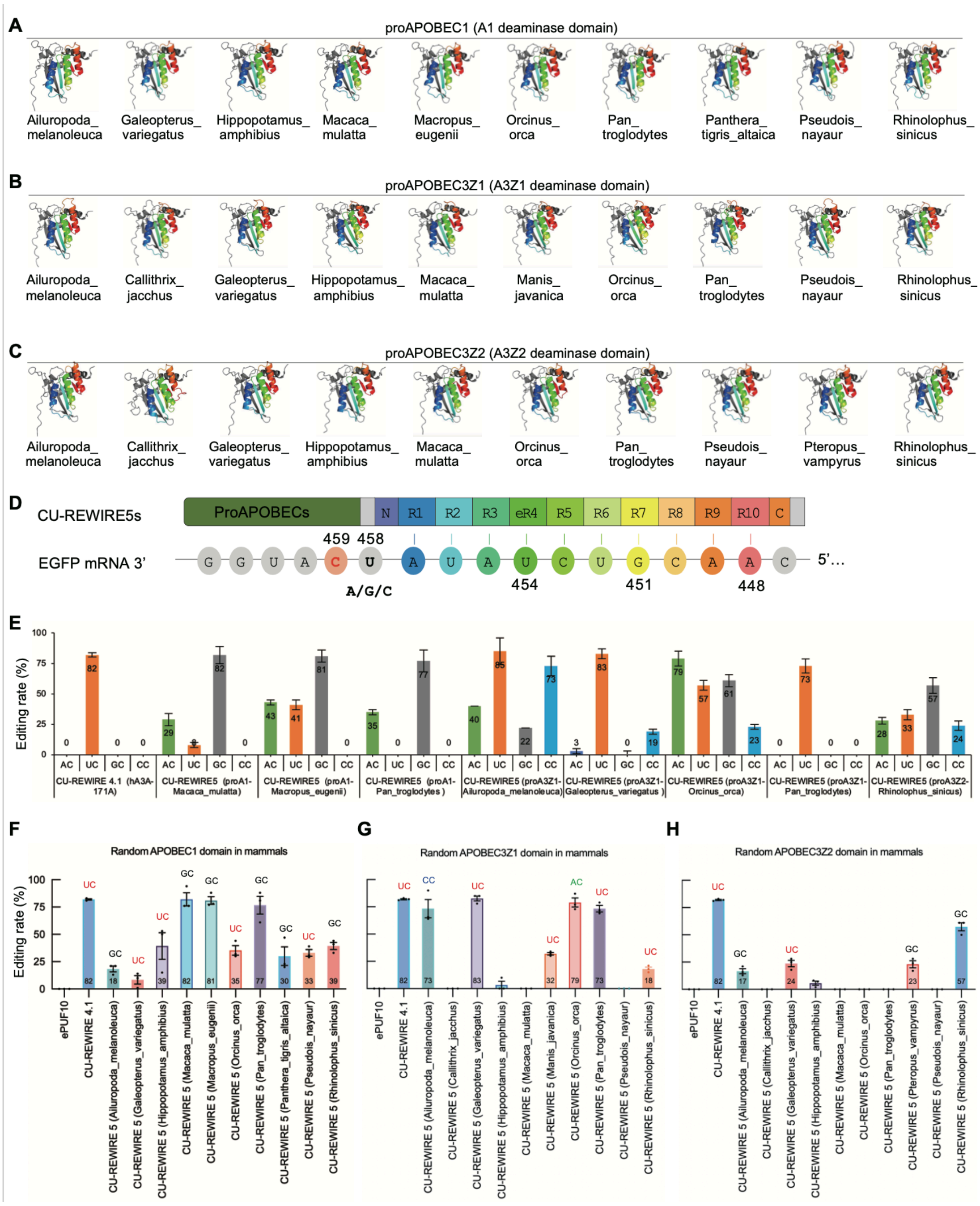


**Figure S5. AI-assisted design of mammal-derived ProAPOBECs for C-to-U editing.**

(A) AlphaFold2-predicted structure of ProAPOBEC1s. The C-to-U deaminase domain of these ProAPOBECs is derived from the APOBEC1 family in mammalian organisms.

(B) AlphaFold2-predicted structure of ProAPOBEC3Z1s. The C-to-U deaminase domain in these ProAPOBECs originates from the APOBEC3 family in mammalian organisms.

(C) AlphaFold2-predicted structure of ProAPOBEC3Z2s. The C-to-U deaminase domain of these ProAPOBECs is also derived from the APOBEC3 family in mammalian organisms.

(D) Illustration of $C_{459}$ recognition in EGFP mRNA by CU-REWIRE5s, as expressed in HEK293T cells.

ePUF10 binding sites in EGFP mRNA and the target $C_{459}$ site are highlighted.

(E) Editing preferences of various CU-REWIREs at the $C_{459}$ site of EGFP mRNA in $A_{458}C$, $U_{458}C$, $G_{458}C$, and $C_{458}C$ contexts.

(F) Efficient editing at the $C_{459}$ site of EGFP mRNA by APOBEC1-derived CU-REWIRE5s. The editing rate of $C_{459}$ was measured by Sanger sequencing. Values represent mean ± SEM (n=3).

(G) Effective editing at the $C_{459}$ site of EGFP mRNA by APOBEC3Z1-derived CU-REWIRE5s.

(H) Efficient editing at the $C_{459}$ site of EGFP mRNA by APOBEC3Z2-derived CU-REWIRE5s.

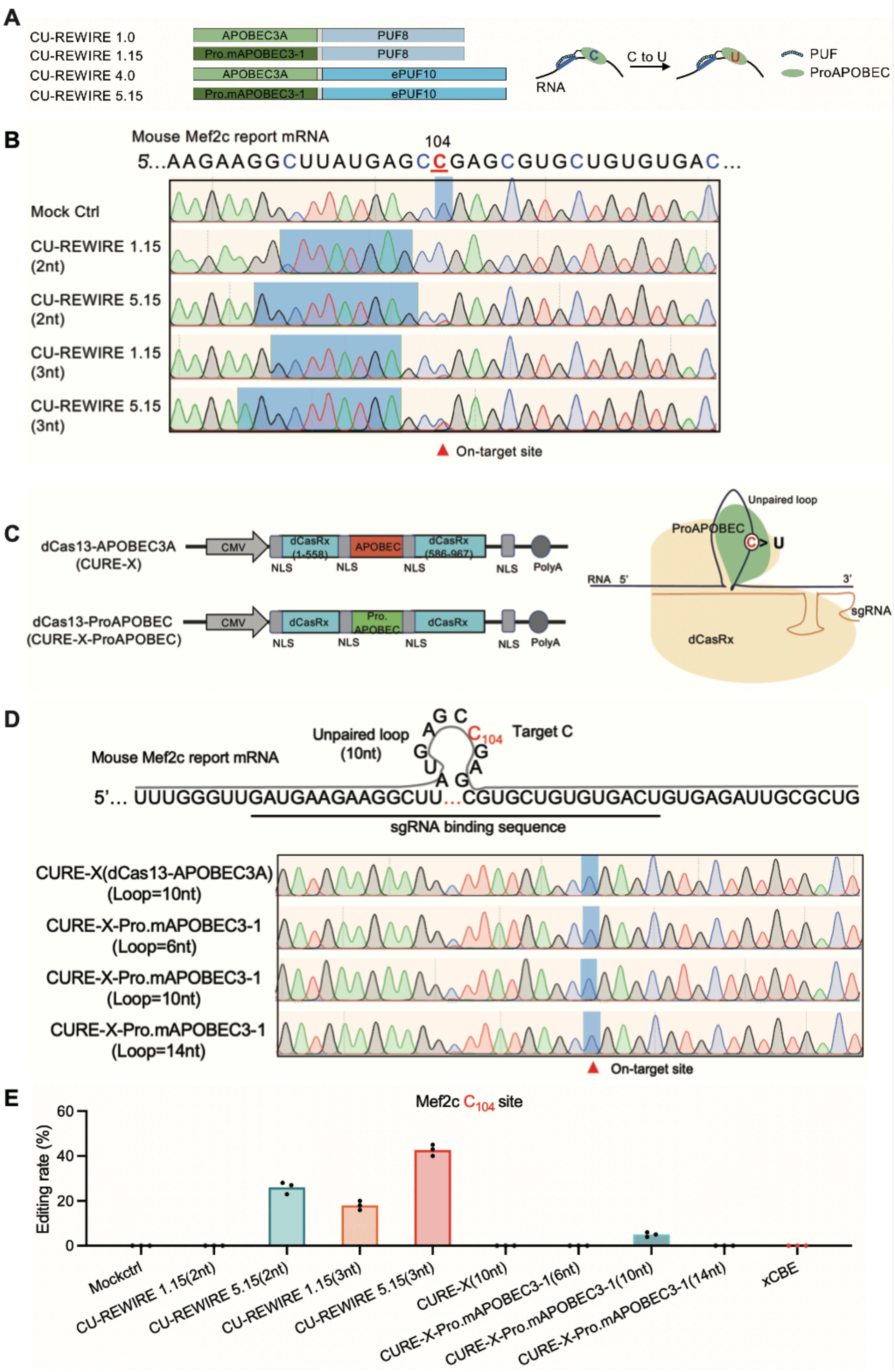


**Figure S6. Editing of mutant Mef2c mRNA by different C-to-U editing tools in HEK293T cell line.**

(A) Schematic of CU-REWIREs with upgraded ProAPOBEC variants. Top, the original CU-REWIRE1s with the PUF8 that recognize targets by 8-nt sequence. Bottom, upgraded CU-REWIRE5s with the ePUF10 that recognize targets by 10-nt sequence.

(B) Efficient editing of $C_{104}$ in Mef2c mRNA by CU-REWIRE1s and CU-REWIRE5s. The PUF binding site in Mef2c is underlined in blue, with on-target site marked in red (top). Editing rates of $C_{104}$ are measured by Sanger-sequencing with triplicates.

(C) Schematic of CURE-X with upgraded ProAPOBEC variants. Top, the original CURE-X fused with the native APOBEC3A. Bottom, upgraded CURE-X fused with the ProAPOBEC.

(D) Efficient editing of $C_{104}$ in Mef2c mRNA by CURE-X-ProAPOBEC. The sgRNA binding sequence in Mef2c is underlined, with on-target site marked in red (top). Editing rates of $C_{104}$ are measured by Sanger-sequencing with triplicates.

(E)Efficient editing of $C_{104}$ in Mef2c mRNA by different C-to-U editing tools. Editing rates of $C_{104}$ are measured by Sanger-sequencing with triplicates.

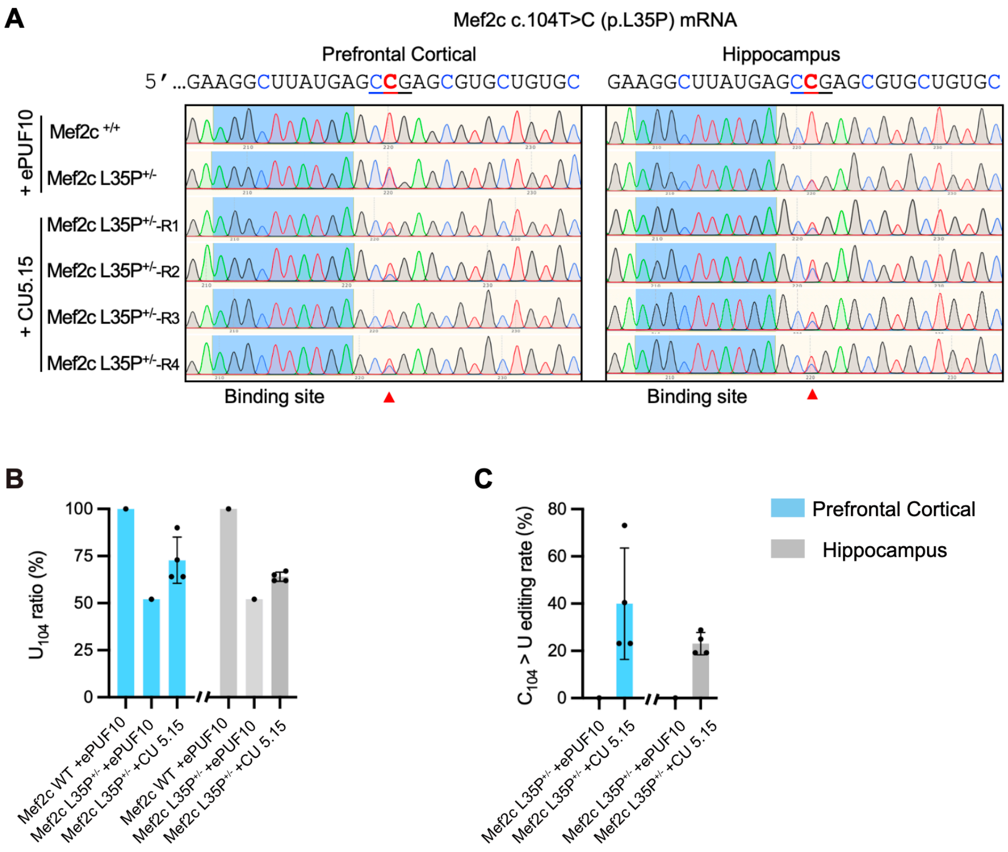


**Figure S7. Effective *in vivo* base editing of $C_{104}$ in Mef2c mutant mice.**

(A) Sanger sequencing results for $C_{104}$ (highlighted in red) in Mef2c mRNA from prefrontal cortical and hippocampal neurons of Mef2c L35P$^{+/-}$ mice, 4-weeks post AAV-CU5.15 injection. Sequencing data shown for 4 independent Mef2c L35P$^{+/-}$ mice (R1-R4), each injected with AAV-CU5.15.

(B) The percentage of $U_{104}$ in prefrontal cortex and hippocampus tissues of Mef2c L35P$^{+/-}$ mice after CU5.15 editing. The editing rates presented were measured by Sanger sequencing.

(C) Efficient editing of Mef2c $C_{104}$ site by CU5.15. The editing rate of $C_{edited}=(U_{CU5.15}-U_{ePUF10}) / C_{total} (1-U_{ePUF10})$.

**Supplementary Tables**

**Table S1. outlines the provenance of ProAPOBECs donors employed in this study.**

| **Current AID/APOBECs deaminase donor in human/mouse/rat** | | | |
|---|---|---|---|
| **CU-REWIREs construction** | **ProAPOBEC name** | **native AID/APOBECs deaminase donor** | **Deaminase domain amino acid sequence (5'-3')** |
| CU-REWIRE5.1 (pro.hA1) | pro.hA1 | human APOBEC1 | HVEVNFIKKFTSERDFHPSMSCSITWFLSWSPCWECSQAIREFLSRHPGVTLVIYVARLFWHMDQQNRQGLRDLVNSGVTIQIM |
| CU-REWIRE5.2 (pro.hAID) | pro.hAID | human AID | HVELLFLRYISDWDLDPGRCYRVTWFTSWSPCYDCARHVADFLRGNPNLSLRIFTARLYFCEDRKAEPEGLRRLHRAGVQIAIM |
| CU-REWIRE5.3 (pro.mA1) | pro.mA1 | mouse APOBEC1 | HVEVNFLEKFTTERYFRPNTRCSITWFLSWSPCGECSRAITEFLSRHPYVTLFIYIARLYHHTDQRNRQGLRDLISSGVTIQIM |
| CU-REWIRE5.4 (pro.mAID) | pro.mAID | mouse AID | HVELLFLRYISDWDLDPGRCYRVTWFTSWSPCYDCARHVAEFLRWNPNLSLRIFTARLYFCEDRKAEPEGLRRLHRAGVQIGIM |
| CU-REWIRE5.5 (pro.hA3B-1) | pro.hA3B-1 | human APOBEC3B | HAELRFLDLVPSLQLDPAQIYRVTWFISWSPCFSWGCAGEVRAFLQENTHVRLRIFAARIYDYDPLYKEALQMLRDAGAQVSIM |
| CU-REWIRE5.6 (pro.hA3B-2) | pro.hA3B-2 | human APOBEC3B | HAEMCFLSWFCGNQLPAYKCFQITWFVSWTPCPDCVAKLAEFLSEHPNVTLTISAARLYYYWERDYRRALCRLSQAGARVKIM |
| CU-REWIRE5.7 (pro.hA3C) | pro.hA3C | human APOBEC3C | HAERCFLSWFCDDILSPNTKYQVTWYTSWSPCPDCAGEVAEFLARHSNVNLTIFTARLYYFQYPCYQEGLRSLSQEGVAVEIM |
| CU-REWIRE5.8 (pro.hA3H) | pro.hA3H | human APOBEC3H | HAEICFINEIKSMGLDETQCYQVTCYLTWSPCSSCAWELVDFIKAHDHLNLGIFASRLYYHWCKPQQKGLRLLCGSQVPVEVM |
| CU-REWIRE5.9 (pro.hA3D-1) | pro.hA3D-1 | human APOBEC3D | HAEMCFLSWFCGNRLPANRRFQITWFVSWNPCLPCVVKVTKFLAEHPNVT |

| | | | |
|---|---|---|---|
| | | | LTISAARLYYYRDRDWRWVLLRLHK AGARVKIM |
| CU-REWIRE5.10 (pro.hA3D-2) | pro.hA3D-2 | human APOBEC3D | HAERCFLSWFCDDILSPNTNYEVTW YTSWSPCPECAGEVAEFLARHSNVN LTIFTARLCYFWDTDYQEGLCSLSQE GASVKIM |
| CU-REWIRE5.11 (pro.hA3F-1) | pro.hA3F-1 | human APOBEC3F | HAEMCFLSWFCGNQLPAYKCFQITW FVSWTPCPDCVAKLAEFLAEHPNVTL TISAARLYYYWERDYRRALCRLSQA GARVKIM |
| CU-REWIRE5.12 (pro.hA3F-2) | pro.hA3F-2 | human APOBEC3F | HAERCFLSWFCDDILSPNTNYEVTW YTSWSPCPECAGEVAEFLARHSNVN LTIFTARLYYFWDTDYQEGLRSLSQE GASVEIM |
| CU-REWIRE5.13 (pro.hA3G-1) | pro.hA3G-1 | human APOBEC3G | HAELCFLDVIPFWKLDLDQDYRVTCF TSWSPCFSCAQEMAKFISKNKHVSLC IFTARIYDDQGRCQEGLRTLAEAGAK ISIM |
| CU-REWIRE5.14 (pro.hA3G-2) | pro.hA3G-2 | human APOBEC3G | HPEMRFFHWFSKWRKLHRDQEYEVT WYISWSPCTKCTRDMATFLAEDPKV TLTIFVARLYYFWDPDYQEALRSLCQ KRDGPRATMKIM |
| CU-REWIRE5.15 (pro.mA3-1) | pro.mA3-1 | mouse APOBEC3 | HAEICFLYWFHDKVLKVLSPREEFKI TWYMSWSPCFECAEQIVRFLATHHN LSLDIFSSRLYNVQDPETQQNLCRLV QEGAQVAAM |
| CU-REWIRE5.16 (pro.mA3-2) | pro.mA3-2 | mouse APOBEC3 | HAEILFLDKIRSMELSQVTITCYLTWS PCPNCAWQLAAFKRDRPDLILHIYTS RLYFHWKRPFQKGLCSLWQSGILVD VM |
| CU-REWIRE5.17 (pro.rA1) | pro.rA1 | rat APOBEC1 | HVEVNFIEKFTTERYFCPNTRCSITWF LSWSPCGECSRAITEFLSRYPHVTLFI YIARLYHHADPRNRQGLRDLISSGVT IQIM |
| **Random A1 deaminase donor in mammals** | | | |
| **CU-REWIREs construction** | **proAPOBEC name** | **native AID/APOBECs deaminase donor** | **Deaminase domain amino acid sequence (5'-3')** |
| CU-REWIRE 5 (proA1- | proA1-Ailuropoda_melanoleuca | A1-Ailuropoda_melanoleuca | HVEINFIEKFTLERQYCPSIHCSVTWFLS WSPCWECSKAIRAFLSQHPSVTLVIYVA RLFWHMEPQNRQGLRDLINSGVTIQIM |

| | | | |
|---|---|---|---|
| Ailuropoda_melanoleuca) | | | |
| CU-REWIRE 5 (proA1-Galeopterus_variegatus) | proA1-Galeopterus_variegatus | A1-Galeopterus_variegatus | HVEVNFIAKFTSERHFHSSISCSITWFLSWSPCWECCKAIREFLSQYPSVTLVIYVARLFQHMDQQNRQGLRDLINSGVTIQII |
| CU-REWIRE 5 (proA1-Hippopotamus_amphibius) | proA1-Hippopotamus_amphibius | A1-Hippopotamus_amphibius | HVERNFIEKITSERHFHPSTSCSIIWFLSWSPCWECSKAIGEFLNQHPSVTLVIYAARLFQHMDPRNRQGLRDLICSGVTIRIM |
| CU-REWIRE 5 (proA1-Macaca_mulatta) | proA1-Macaca_mulatta | A1-Macaca_mulatta | HVEVNFIEKLTSERRFHSSISCSITWFLSWSPCWECSQAIREFLSQHPGVTLVIYVARLFWHTDQQNRQGLRDLVNSGVTIQIM |
| CU-REWIRE 5 (proA1-Macropus_eugenii) | proA1-Macropus_eugenii | A1-Macropus_eugenii | HAEINFMEKFTSERHFKSSVKCSITWFLSWSPCWRCSKAIREFLNQHPNVTLIIFVARLYWHMDQRHRQELKELHSSGVAIWIM |
| CU-REWIRE 5 (proA1-Orcinus_orca) | proA1-Orcinus_orca | A1-Orcinus_orca | HVECNFIEKVTSERSFHRSVSCCIIWFLSWSPCWECSKAIREFLNQHPRVTLRIYVARLFQHMDRRNRQGLRDLIRSGVTIQIM |
| CU-REWIRE 5 (proA1-Pan_troglodytes) | proA1-Pan_troglodytes | A1-Pan_troglodytes | HVEVNFIKKFTSERHFHPSISCSITWFLSWSPCWECSQAIREFLSQHPGVTLVIYVARLFWHMDQQNRQGLRDLVNSGVTIQIM |
| CU-REWIRE 5 (proA1-Panthera_tigris_altaica) | proA1-Panthera_tigris_altaica | A1-Panthera_tigris_altaica | HVELNFIEKFTSERHFCPSVSCSITWFLSWSPCWECSKAIRGFLSQHPSVTLVIYVSRLFWHLDQQNRQGLRDLINSGVTVQIM |
| CU-REWIRE 5 (proA1-Pseudois_nayaur) | proA1-Pseudois_nayaur | A1-Pseudois_nayaur | HVERNFIEKIASERHFRPSISCSIFWYLSWSPCWECSKAIREFLNQHPNVTLVIYIARLFQHTDPQNRQGLKDLFHSGVTIQVM |
| CU-REWIRE 5 (proA1-Rhinolophus_sinicus) | proA1-Rhinolophus_sinicus | APOBEC1-Rhinolophus_sinicus | HVELNFIEKFTSERRFCSSISCSIIWFLSWSPCWECSKAIREFLSQRPGVTLVIYAARLYWHMDQQNRQGLRDLINCGVTIQIM |
| **Random A3Z1 deaminase donor in mammals** | | | |
| **CU-REWIREs construction** | **proAPOBEC name** | **native AID/APOBECs deaminase donor** | **Deaminase domain amino acid sequence (5’-3’)** |
| CU-REWIRE 5 (proA3Z1-Ailuropoda_melanoleuca) | proA3Z1-Ailuropoda_melanoleuca | A3Z1-Ailuropoda_melanoleuca | HAECYLLEQIQSWNLDPKLHYGVTCFLSWSPCAKCAQKMARFLQENSHVSLKLFASRLYTRERWDEDYKEGLRTLKRAGASIAIM |

| | | | |
|---|---|---|---|
| CU-REWIRE 5 (proA3Z1-Callithrix_jacchus) | proA3Z1-Callithrix_jacchus | A3Z1-Callithrix_jacchus | HAELCFLDLISFWKLNLAQPYRVTCFISWSTCFSCAQMMAKFLQENTHVSLSTFADLICDHHGRYKEGLRRLDRAGAPISMI |
| CU-REWIRE 5 (proA3Z1-Galeopterus_variegatus) | proA3Z1-Galeopterus_variegatus | A3Z1-Galeopterus_variegatus | HAELCFLDLLNSWQLYPALRYRVTWFISWSPCCNCAQEVAAFLGWNSHVHLRIFAARIHDRFPGYEQGLRTLQGAGAQISIM |
| CU-REWIRE 5 (proA3Z1-Hippopotamus_amphibius) | proA3Z1-Hippopotamus_amphibius | A3Z1-Hippopotamus_amphibius | HAELFFLDRIRSWNLDRRLLYRLTCFISWTPCHTCAQKLAMFLRENSHVSLNMFASRIYSLNDYEAGLRTLQAAGAQINIM |
| CU-REWIRE 5 (proA3Z1-Macaca_mulatta) | proA3Z1-Macaca_mulatta | A3Z1-Macaca_mulatta | HVEQCFLDLVSSWHLDPAQCYRIIYFISWSSCFNCAQEIAGFLRENRHMSLHIFTACIYDYHPGYEERLHMLQGIGAQISIM |
| CU-REWIRE 5 (proA3Z1-Manis_eugenii) | proA3Z1-Macropus_eugenii | A3Z1-Macropus_eugenii | HAEFLLLERILSWKLNPKLRYMVNCFISWSPCAACAQYIANFLRENLHVHLYIFASRIYRRNDYKEGLRTVWEAGAQMAIM |
| CU-REWIRE 5 (proA3Z1-Orcinus_orca) | proA3Z1-Orcinus_orca | A3Z1-Orcinus_orca | HAEHYFLDRIRSWNLDRELHYRLTCFISWTPCDTCAQRLADFLGKNSHVSLHIFASRIYSLSDYEAGLRTLQAAGAQIAIM |
| CU-REWIRE 5 (proA3Z1-Pan_troglodytes) | proA3Z1-Pan_troglodytes | A3Z1-Pan_troglodytes | HAELRFLDLVPSLQLDPAQIYRVTWFISWSPCFSWGCAGQVRAFLQENTHVRLRIFAARIYDYDPLYKEALQMLRDAGAQVSIM |
| CU-REWIRE 5 (proA3Z1-Pseudois_nayaur) | proA3Z1-Pseudois_nayaur | A3Z1-Pseudois_nayaur | HAELYFLEQIRSWNLDRDQRYRLTCFISWTPCYNCAQKLTTFLEENRHISLHIFASRIYSVKDSGCHRSGLRALQERARITIM |
| CU-REWIRE 5 (proA3Z1-Rhinolophus_sinicus) | proA3Z1-Rhinolophus_sinicus | A3Z1-Rhinolophus_sinicus | HAELCLLRLIRSWELNTEQHYRVTCFISWSPCHDCARELAAFLGKNSHLSLCVFASHIYTLGGYKTGLRQLQAAGAQVAIM |
| **Random A3Z2 deaminase donor in mammals** | | | |
| **CU-REWIREs construction** | **proAPOBEC name** | **native AID/APOBECs deaminase donor** | **Deaminase domain amino acid sequence (5’-3’)** |
| CU-REWIRE 5 (proA3Z2-Ailuropoda_melanoleuca) | proA3Z2-Ailuropoda_melanoleuca | A3Z2-Ailuropoda_melanoleuca | HAESCFLSWFRAQNLSPDEDYHVTWFSSWSPCHTCADEVVEFLGQYRHVTLSIFAARLYYFWDPPFQNGLRRLQSAGVRLDIM |
| CU-REWIRE 5 (proA3Z2-Callithrix_jacchus) | proA3Z2-Callithrix_jacchus | A3Z2-Callithrix_jacchus | HAEMRFLHWFRKWKLYSDQEYEVTWFVSWSPCPVCARNVAEFLARDRKVTLTIFVA |

| | | | |
|---|---|---|---|
| | | | RLYYFWDPHYREELRRLCQKKKNPHATMKIM |
| CU-REWIRE 5 (proA3Z2-Galeopterus_variegatus) | proA3Z2-Galeopterus_variegatus | A3Z2-Galeopterus_variegatus | HAELCFLSWFFNNVLSHDKHYLVTWYISWSPCCKCAEQVADFLARHRNVTLTIFTARLYYFWESEYRQGLRRLCWEGAQVNVM |
| CU-REWIRE 5 (proA3Z2-Hippopotamus_amphibius) | proA3Z2-Hippopotamus_amphibius | A3Z2-Hippopotamus_amphibius | HAEISFLSWFRAEMLSPDEQYEVTWFLSWSPCLACAEQVAEFLRQNRNVTLRIFAARLYYFWKPEYRAWLRRLHHQGTLVNIM |
| CU-REWIRE 5 (proA3Z2-Macaca_mulatta) | proA3Z2-Macaca_mulatta | A3Z2-Macaca_mulatta | HAEMCFLSWFCGNQLPAYKRFQITWFVSWNPCPDCVAKVTEFLAEHPNVTLTISAARLYYYWGKDWRRALRRLHQAGARVKIM |
| CU-REWIRE 5 (proA3Z2-Orcinus_orca) | proA3Z2-Orcinus_orca | A3Z2-Orcinus_orca | HVELCFLSWFRAKKLSSHEQYRITWFLSWSPCLSCAEQVVAFLKENRNVRLSIFAARLYYFWKPDYQHGLRILRHQRAWVRIM |
| CU-REWIRE 5 (proA3Z2-Pan_troglodytes) | proA3Z2-Pan_troglodytes | A3Z2-Pan_troglodytes | HAEMCFLSWFCGNQLSAYKCFQITWFVSWTPCPDCVAKLAKFLAEHPNVTLTISAARLYYYWERDYRRALCRLSQAGARVKIM |
| CU-REWIRE 5 (proA3Z2-Pteropus_vampyrus) | proA3Z2-Pteropus_vampyrus | A3Z2-Pteropus_vampyrus | HAELCFLNWFRAEKLSPYEHYDVTWFLSWSPCSTCAKKIAIFLSNHKNVRLSIFVSRIYYFWKPAFRQGLQELDHLGVQLDAM |
| CU-REWIRE 5 (proA3Z2-Pseudois_nayaur) | proA3Z2-Pseudois_nayaur | A3Z2-Pseudois_nayaur | HSERRFLSWFCAKELRPDECYHITWFMSWSPCMKCAELVAGFLGMYQNVTLSIFAARLYYFQKPQYRMGLLGLSDQGARVDIM |
| CU-REWIRE 5 (proA3Z2-Rhinolophus_sinicus) | proA3Z2-Rhinolophus_sinicus | A3Z2-Rhinolophus_sinicus | HAELCFLSWFKANKLSPGEHYHVTWFLSWSPCARCAEEIANFLKQHRNVTLRVFAARLYYFWNPAFQQGLRKLRCLGAQVKVM |

#### Table S2. Summary of RNA-seq.

| **Biosample_accession of RNA-seq data** | | | |
|---|---|---|---|
| **Sample name _HEK293T** | **Biosample _accession** | **Library _selection** | **Mapped reads** |
| CU-REWIRE3.0(hA3A-PUF10)_EGFP-UC_1 | SAMN39596128 | mRNA-seq | 20M |
| CU-REWIRE3.0(hA3A-PUF10)_EGFP-UC_2 | SAMN39596129 | mRNA-seq | 20M |
| CU-REWIRE3.0(hA3A-PUF10)_EGFP-UC_3 | SAMN39596130 | mRNA-seq | 20M |
| CU-REWIRE4.0(hA3A-ePUF10)_EGFP-WT_1 | SAMN39596131 | mRNA-seq | 20M |

| | | | |
|---|---|---|---|
| CU-REWIRE4.0(hA3A-ePUF10)_EGFP-WT_2 | SAMN39596132 | mRNA-seq | 20M |
| CU-REWIRE4.0(hA3A-ePUF10)_EGFP-WT_3 | SAMN39596133 | mRNA-seq | 20M |
| HEK293T_EGFP-Mock_1 | SAMN39596134 | mRNA-seq | 20M |
| HEK293T_EGFP-Mock_2 | SAMN39596135 | mRNA-seq | 20M |
| HEK293T_EGFP-Mock_3 | SAMN39596136 | mRNA-seq | 20M |
| HEK293T_EGFP-Mock_4 | SAMN39596137 | mRNA-seq | 20M |
| HEK293T_PUF10-Ctrl_1 | SAMN39596138 | mRNA-seq | 20M |
| HEK293T_PUF10-Ctrl_2 | SAMN39596139 | mRNA-seq | 20M |
| HEK293T_PUF10-Ctrl_3 | SAMN39596140 | mRNA-seq | 20M |
| HEK293T_PUF10-Ctrl_4 | SAMN39596141 | mRNA-seq | 20M |
| CU-REWIRE4.1(hA3A-C171A-ePUF10)_EGFP-WT_1 | SAMN39596142 | mRNA-seq | 20M |
| CU-REWIRE4.1(hA3A-C171A-ePUF10)_EGFP-WT_2 | SAMN39596143 | mRNA-seq | 20M |
| CU-REWIRE4.1(hA3A-C171A-ePUF10)_EGFP-WT_3 | SAMN39596144 | mRNA-seq | 20M |
| CU-REWIRE4.2(hA3A-H11A-171A-ePUF10)_EGFP-WT_1 | SAMN39596145 | mRNA-seq | 20M |
| CU-REWIRE4.2(hA3A-H11A-171A-ePUF10)_EGFP-WT_2 | SAMN39596146 | mRNA-seq | 20M |
| CU-REWIRE4.2(hA3A-H11A-171A-ePUF10)_EGFP-WT_3 | SAMN39596147 | mRNA-seq | 20M |
| CU-REWIRE4.3(hA3A-H16A-171A-ePUF10)_EGFP-WT_1 | SAMN39596148 | mRNA-seq | 20M |
| CU-REWIRE4.3(hA3A-H16A-171A-ePUF10)_EGFP-WT_2 | SAMN39596149 | mRNA-seq | 20M |
| CU-REWIRE4.3(hA3A-H16A-171A-ePUF10)_EGFP-WT_3 | SAMN39596150 | mRNA-seq | 20M |
| CU-REWIRE4.5(hA3A-H56A-171A-ePUF10)_EGFP-WT_1 | SAMN39596151 | mRNA-seq | 20M |
| CU-REWIRE4.5(hA3A-H56A-171A-ePUF10)_EGFP-WT_2 | SAMN39596152 | mRNA-seq | 20M |
| CU-REWIRE4.5(hA3A-H56A-171A-ePUF10)_EGFP-WT_3 | SAMN39596153 | mRNA-seq | 20M |

| | | | |
|---|---|---|---|
| CU-REWIRE4.6(hA3A-11A-16A-171A-ePUF10)_EGFP-WT_1 | SAMN39596154 | mRNA-seq | 20M |
| CU-REWIRE4.6(hA3A-11A-16A-171A-ePUF10)_EGFP-WT_2 | SAMN39596155 | mRNA-seq | 20M |
| CU-REWIRE4.6(hA3A-11A-16A-171A-ePUF10)_EGFP-WT_3 | SAMN39596156 | mRNA-seq | 20M |
| CU-REWIRE4.7(hA3A-11A-56A-171A-ePUF10)_EGFP-WT_1 | SAMN39596157 | mRNA-seq | 20M |
| CU-REWIRE4.7(hA3A-11A-56A-171A-ePUF10)_EGFP-WT_2 | SAMN39596158 | mRNA-seq | 20M |
| CU-REWIRE4.7(hA3A-11A-56A-171A-ePUF10)_EGFP-WT_3 | SAMN39596159 | mRNA-seq | 20M |
| CU-REWIRE4.8(hA3A-16A-56A-171A-ePUF10)_EGFP-WT_1 | SAMN39596160 | mRNA-seq | 20M |
| CU-REWIRE4.8(hA3A-16A-56A-171A-ePUF10)_EGFP-WT_2 | SAMN39596161 | mRNA-seq | 20M |
| CU-REWIRE4.8(hA3A-16A-56A-171A-ePUF10)_EGFP-WT_3 | SAMN39596162 | mRNA-seq | 20M |
| CU-REWIRE4.9(hA3A-11A-16A-56A-171A-ePUF10)_EGFP-WT_1 | SAMN39596163 | mRNA-seq | 20M |
| CU-REWIRE4.9(hA3A-11A-16A-56A-171A-ePUF10)_EGFP-WT_2 | SAMN39596164 | mRNA-seq | 20M |
| CU-REWIRE4.9(hA3A-11A-16A-56A-171A-ePUF10)_EGFP-WT_3 | SAMN39596165 | mRNA-seq | 20M |
| CU-REWIRE5.1(Pro.hAPOBEC1-ePUF10)_EGFP-GC_1 | SAMN39596166 | mRNA-seq | 20M |
| CU-REWIRE5.1(Pro.hAPOBEC1-ePUF10)_EGFP-GC_2 | SAMN39596167 | mRNA-seq | 20M |
| CU-REWIRE5.1(Pro.hAPOBEC1-ePUF10)_EGFP-GC_3 | SAMN39596168 | mRNA-seq | 20M |
| CU-REWIRE5.3(Pro.mAPOBEC1-ePUF10)_EGFP-UC_1 | SAMN39596169 | mRNA-seq | 20M |
| CU-REWIRE5.3(Pro.mAPOBEC1-ePUF10)_EGFP-UC_2 | SAMN39596170 | mRNA-seq | 20M |
| CU-REWIRE5.3(Pro.mAPOBEC1-ePUF10)_EGFP-UC_3 | SAMN39596171 | mRNA-seq | 20M |
| CU-REWIRE5.7(Pro.hAPOBEC3C-ePUF10)_EGFP-UC_1 | SAMN39596172 | mRNA-seq | 20M |
| CU-REWIRE5.7(Pro.hAPOBEC3C-ePUF10)_EGFP-UC_2 | SAMN39596173 | mRNA-seq | 20M |
| CU-REWIRE5.7(Pro.hAPOBEC3C-ePUF10)_EGFP-UC_3 | SAMN39596174 | mRNA-seq | 20M |
| CU-REWIRE5.8(Pro.hAPOBEC3H-ePUF10)_EGFP-GC_1 | SAMN39596175 | mRNA-seq | 20M |

| | | | |
|---|---|---|---|
| CU-REWIRE5.8(Pro.hAPOBEC3H-ePUF10)_EGFP-GC_2 | SAMN39596176 | mRNA-seq | 20M |
| CU-REWIRE5.8(Pro.hAPOBEC3H-ePUF10)_EGFP-GC_3 | SAMN39596177 | mRNA-seq | 20M |
| CU-REWIRE5.13(Pro.hAPOBEC3G-1-ePUF10)_EGFP-UC_1 | SAMN39596178 | mRNA-seq | 20M |
| CU-REWIRE5.13(Pro.hAPOBEC3G-1-ePUF10)_EGFP-UC_2 | SAMN39596179 | mRNA-seq | 20M |
| CU-REWIRE5.13(Pro.hAPOBEC3G-1-ePUF10)_EGFP-UC_3 | SAMN39596180 | mRNA-seq | 20M |
| CU-REWIRE5.15(Pro.mAPOBEC3-1-ePUF10)_EGFP-CC_1 | SAMN39596181 | mRNA-seq | 20M |
| CU-REWIRE5.15(Pro.mAPOBEC3-1-ePUF10)_EGFP-CC_2 | SAMN39596182 | mRNA-seq | 20M |
| CU-REWIRE5.15(Pro.mAPOBEC3-1-ePUF10)_EGFP-CC_3 | SAMN39596183 | mRNA-seq | 20M |
| CU-REWIRE5.16(Pro.mAPOBEC3-2-ePUF10)_EGFP-AC_1 | SAMN39596184 | mRNA-seq | 20M |
| CU-REWIRE5.16(Pro.mAPOBEC3-2-ePUF10)_EGFP-AC_2 | SAMN39596185 | mRNA-seq | 20M |
| CU-REWIRE5.16(Pro.mAPOBEC3-2-ePUF10)_EGFP-AC_3 | SAMN39596186 | mRNA-seq | 20M |
| CU-REWIRE5.17(Pro.rAPOBEC1-ePUF10)_EGFP-GC_1 | SAMN39596187 | mRNA-seq | 20M |
| CU-REWIRE5.17(Pro.rAPOBEC1-ePUF10)_EGFP-GC_2 | SAMN39596188 | mRNA-seq | 20M |
| CU-REWIRE5.17(Pro.rAPOBEC1-ePUF10)_EGFP-GC_3 | SAMN39596189 | mRNA-seq | 20M |
| **Sample name _mouse** | **Biosample _accession** | **Library _selection** | **Mapped reads** |
| Prefrontal Cortex_Mef2c WT_Ctrl_1 | SAMN39608906 | mRNA-seq | 20M |
| Prefrontal Cortex_Mef2c WT_Ctrl_2 | SAMN39608907 | mRNA-seq | 20M |
| Prefrontal Cortex_Mef2c WT_Ctrl_3 | SAMN39608908 | mRNA-seq | 20M |
| Prefrontal Cortex_Mef2c WT_PUF10-Ctrl_1 | SAMN39608909 | mRNA-seq | 20M |
| Prefrontal Cortex_Mef2c WT_PUF10-Ctrl_2 | SAMN39608910 | mRNA-seq | 20M |
| Prefrontal Cortex_Mef2c WT_PUF10-Ctrl_3 | SAMN39608911 | mRNA-seq | 20M |
| Prefrontal Cortex_Mef2c L35P+/-_PUF10-Ctrl_1 | SAMN39608912 | mRNA-seq | 20M |

| | | | |
|---|---|---|---|
| Prefrontal Cortex_Mef2c L35P+/-_PUF10-Ctrl_2 | SAMN39608913 | mRNA-seq | 20M |
| MPrefrontal Cortex_ef2c L35P+/-_PUF10-Ctrl_3 | SAMN39608914 | mRNA-seq | 20M |
| Prefrontal Cortex_Mef2c L35P+/-_CU-REWIRE5.15_1 | SAMN39608915 | mRNA-seq | 20M |
| Prefrontal Cortex_Mef2c L35P+/-_CU-REWIRE5.15_2 | SAMN39608916 | mRNA-seq | 20M |
| Prefrontal Cortex_Mef2c L35P+/-_CU-REWIRE5.15_3 | SAMN39608917 | mRNA-seq | 20M |
| Hippocampus_Mef2c WT_Ctrl_1 | SAMN39608918 | mRNA-seq | 20M |
| Hippocampus_Mef2c WT_Ctrl_2 | SAMN39608919 | mRNA-seq | 20M |
| Hippocampus_Mef2c WT_Ctrl_3 | SAMN39608920 | mRNA-seq | 20M |
| Hippocampus_Mef2c WT_PUF10-Ctrl_1 | SAMN39608921 | mRNA-seq | 20M |
| Hippocampus_Mef2c WT_PUF10-Ctrl_2 | SAMN39608922 | mRNA-seq | 20M |
| Hippocampus_Mef2c WT_PUF10-Ctrl_3 | SAMN39608923 | mRNA-seq | 20M |
| Hippocampus_Mef2c L35P+/-_PUF10-Ctrl_1 | SAMN39608924 | mRNA-seq | 20M |
| Hippocampus_Mef2c L35P+/-_PUF10-Ctrl_2 | SAMN39608925 | mRNA-seq | 20M |
| Hippocampus_Mef2c L35P+/-_PUF10-Ctrl_3 | SAMN39608926 | mRNA-seq | 20M |
| Hippocampus_Mef2c L35P+/-_CU-REWIRE5.15_1 | SAMN39608927 | mRNA-seq | 20M |
| Hippocampus_Mef2c L35P+/-_CU-REWIRE5.15_2 | SAMN39608928 | mRNA-seq | 20M |
| Hippocampus_Mef2c L35P+/-_CU-REWIRE5.15_3 | SAMN39608929 | mRNA-seq | 20M |
